# Integrative Omics and Network Biology Reveal Transcriptional Changes of Amino Acid Transport in Arabidopsis Susceptibility to *Pseudomonas syringae*

**DOI:** 10.64898/2026.03.25.714176

**Authors:** Bharat Mishra, Nilesh Kumar, Yali Sun, Thomas W. Detchemendy, Doni Thingujam, Adrian Flannery, Karolina M. Pajerowska-Mukhtar, M. Shahid Mukhtar

## Abstract

Plant amino acids function as both pathogen nutrients and essential drivers of systemic immunity. The regulation of amino acid homeostasis through transporters is a essential for mounting a robust and coordinated immune response in plants during pathogen infection. Using systems biology and integrative network science, we investigated bacterial virulence in Arabidopsis. By comparing gene coexpression networks of effector-triggered susceptibility (ETS) and pattern-triggered immunity (PTI), we uncovered a plant amino acid-related processes specifically linked to ETS. Integrating time-series transcriptomics, protein–DNA interactions, and mathematical simulations, we identified ANAC046 as a transcriptional regulator of amino acid processes during ETS. Single-cell RNA-Seq revealed that amino acid transporters are primarily expressed in companion and mesophyll cells, while functional validation confirmed ANAC046’s roles in promoting susceptibility. Further integration of transcriptome and interactome data showed that amino acid-related genes interact with key immune hub proteins. Network topology analysis enabled the characterization of seven additional genes involved in plant defense. To support community-wide research, we developed MID*ata*, an open-access platform for pre-analyzed Arabidopsis networks. Together, our findings demonstrate the power of systems-level approaches in uncovering hierarchical regulatory mechanisms underlying plant susceptibility to bacterial pathogens.

## Introduction

Static and dynamic molecular networks orchestrate a multitude of regulatory pathways and signaling cascades to respond to diverse environmental perturbations, physiological signals, and pathogen infections (Mukhtar 2013, Mukhtar, McCormack et al. 2016, Lopez and Mukhtar 2017, Gonzalez-Fuente, Carrere et al. 2020). These networks are based on assemblages of protein-protein, protein-nucleic acids, and protein-ligand complexes. The growing availability of large-scale -omics datasets including those for sequencing of nucleic acids (genomics and transcriptomics), protein-nucleic acid interactions (gene regulatory relationships), quantitative measurements of proteins (proteomics), detection of protein-protein interactions (PPIs) at a global scale (interactomics), and organismal changes in phenotype due to internal and external signals (phenomics) is driving the emergence of systems-levels approaches (Garbutt, Bangalore et al. 2014, Mishra, Sun et al. 2017, Mishra, Sun et al. 2018, Gordon, Hiatt et al. 2020, Khorsand, Savadi et al. 2020). Systems biology entails generating, integrating, analyzing, modeling, simulating, and inferring datasets from diverse experimental -omics sources (Garbutt, Bangalore et al. 2014). Network science, a branch of systems biology, represents, analyzes, and visualizes biological systems using graph theory tools, and ultimately interprets the complexities of molecular interactions into biological messages (Garbutt, Bangalore et al. 2014, Mishra, Kumar et al. 2019). Network biology describes molecular interactions as graphs composed of vertices (nodes) and edges (links or connections among vertices). In a biological network, genes and their products constitute nodes while functional or physical relationships among these nodes constitute edges. These networks—built from gene coexpression, protein–protein interactions, or transcriptional regulation—serve as a framework to explore how cellular processes are coordinated, reprogrammed, or perturbed over time and across different spatial contexts(Kumar and Mukhtar 2023, Kumar and Mukhtar 2023, Thingujam, Tan et al. 2025). Functional modules are embedded within these biological systems and can execute diverse cellular functions (Garbutt, Bangalore et al. 2014, Mishra, Kumar et al. 2019). Network-based approaches are highly effective for uncovering key regulatory components—often referred to as hubs or bottlenecks—that control biological systems(Kumar and Mukhtar 2023, Kumar and Mukhtar 2023). By applying topological analysis and comparative modeling, network biology helps reveal molecular signatures associated with distinct cellular states, providing a powerful framework for understanding both steady-state cellular and physiological processes, or disease mechanisms. Weighted Gene Coexpression Network Analysis (WGCNA)(Langfelder and Horvath 2008), for example, has emerged as a widely adopted method to construct gene coexpression modules and reveal key genes that mediate transitions between cellular states. Integrating temporal, single-cell and spatial, and context-specific data further refines these insights, enabling a mechanistic understanding of dynamic responses to environmental cues or biotic stressors(Mishra, Sun et al. 2017, Zhu, Lolle et al. 2023, Thingujam, Tan et al. 2025). While small networks and pathways are easy to store and interpret, complex networks necessitate new tools derived from graph theory. System and network biology perspectives have provided important insights into many biological processes but are only just beginning to be applied to plant-pathogen interactions (Mishra, Kumar et al. 2019). Such comprehensive models hold promise in deciphering complex interactions, especially in the context of plant–microbe relationships, where multiple layers of regulation and signaling are at play.

Plant-microbe interactions are multifaceted, involving a finely tuned balance between immune responses and microbial strategies to evade or suppress these defenses(Zhu, Moreno-Perez et al. 2023). From a systems biology perspective, these interactions are best understood not as binary outcomes of resistance versus susceptibility, but as dynamic regulatory programs shaped by coevolving molecular interactions(Mishra, Kumar et al. 2019, Mishra, Kumar et al. 2021). Plants can detect both conserved molecular patterns and secreted virulence proteins (effectors) to activate patterns- and effector-triggered immunity (PTI and ETI, respectively) (Mukhtar 2013, Pajerowska-Mukhtar, Emerine et al. 2013, Lopez, Sun et al. 2015, Smakowska-Luzan, Mott et al. 2018). Pathogens deploy suites of effector proteins and RNAs to perturb or manipulate diverse plant functional modules, including protein-DNA and protein-protein clusters to promote pathogen colonization. The effects of these manipulations are commonly referred to as effector-triggered susceptibility (ETS) (Washington, Mukhtar et al. 2016, Sun, Detchemendy et al. 2018, Zaidi, Mukhtar et al. 2018). Despite significant progress in identifying individual genes and signaling pathways involved in plant defense, the full complexity of transcriptional reprogramming during pathogen infection remains poorly understood. Transcriptional responses are not merely additive but arise from complex regulatory cascades, feedback loops, and combinatorial control by transcription factors (TFs) and chromatin modifiers(Kim, Kidokoro et al. 2024, Shu, Li et al. 2024, Beckers, Mamiya et al. 2025). Network biology offers a framework to model these responses at a systems level, identify condition-specific coexpression modules, and predict regulatory hubs that orchestrate context-dependent gene expression(Mishra, Kumar et al. 2021, Kumar and Mukhtar 2023). In the context of plant-pathogen interactions, constructing and comparing gene coexpression networks under PTI- and ETS-inducing conditions allows for the identification of shared and unique regulatory circuits that define resistance or susceptibility states. A key feature of ETS is the suppression of host immune responses, achieved when pathogen effectors specifically target critical immune regulatory hubs (Mishra, Kumar et al. 2021, Kumar and Mukhtar 2023). A particularly underexplored aspect of plant-microbe interactions is the role of host metabolism, especially amino acid biosynthesis and transport, in determining the outcome of infection(Yao, Sui et al. 2025). Pathogens are known to reprogram host metabolism to meet their nutritional demands or to evade defense(Sonawala, Dinkeloo et al. 2018, Wang, Liu et al. 2022). Recent studies have suggested that metabolic pathways are not merely passive consequences of infection but are actively modulated to favor pathogen proliferation(Sonawala, Dinkeloo et al. 2018, Wang, Liu et al. 2022, Yao, Sui et al. 2025). Additionally, recent advances in single-cell RNA sequencing (scRNA-Seq) technologies provide unprecedented resolution into the cellular specificity of gene expression during infection (Zhu, Lolle et al. 2023, Zhu, Moreno-Perez et al. 2023, Nobori, Monell et al. 2025, Thingujam, Tan et al. 2025). It offers the opportunity to pinpoint specific cell types where susceptibility factors—such as amino acid transporters and their regulatory transcription factors—are most active. Such insights could inform tissue-specific strategies for enhancing resistance. Integrative network biology enables the identification of key pathways by detecting co-regulated modules within specific cell-types enriched in metabolic functions and linking them to upstream transcriptional regulators.

While previous studies provide insight on how pathogens exploit host nutrient transport systems, relatively little is known about the regulatory mechanisms governing amino acid transporters in plants during infection. One of the few known examples is bZIP11, a transcription factor shown to regulate *UMAMIT* (usually multiple acids move in and out transporters) and *SWEET* (Sugars Will Eventually Be Exported Transporters) genes that facilitate nutrient efflux from host cells—processes that can be hijacked by pathogens like *Pseudomonas syringae* pv. tomato DC3000 (hereafter DC3000) to support their proliferation(Prior, Selvanayagam et al. 2021). But beyond bZIP11, how a broader array of other amino acid transporters such as amino acid permeases (AAPs), lysine histidine transporters (LHTs), amino acid transporters (AATs) families or other amino acid-related genes is transcriptionally controlled has remained largely unexplored. In this study, we sought to comprehensively characterize the transcriptional networks and regulatory logic that underlie ETS in Arabidopsis in response to a bacterial pathogen hereafter DC3000. Towards this, we conducted a comparative coexpression network analysis on transcriptome datasets from infections with DC3000 and DC3000 *hrpA*^−^ Type III secretion defective mutant (hereafter hrpA^−^), revealing that the ETS-specific network is uniquely enriched in amino acid metabolism pathways. To elucidate the mechanistic basis of transcriptional rewiring, we integrated temporal gene expression with static TF-target networks using probabilistic graphical models, which identified 85 ETS-specific transcription factors. Among these, ANAC046 emerged as a master regulator orchestrating amino acid-associated gene expression. Single-cell RNA-Seq analysis further revealed that amino acid transporter genes were highly expressed in companion and mesophyll cells, and ANAC046 expression was localized to mesophyll and vascular cells. Functional assays using Arabidopsis plants with reduced ANAC046 expression confirmed that amino acid-related genes depend on ANAC046 and supported its role in ETS, as these plants showed increased resistance specifically to DC3000 and elevated expression of salicylic acid-responsive genes. By integrating bulk and single-cell transcriptomics with protein–protein interaction networks and network topology analysis, we revealed that amino acid metabolism genes interact with hub proteins. Furthermore, through topological analysis—examining various centrality metrics—we identified seven additional genes with previously uncharacterized roles in plant defense. Finally, we developed MID*ata*, an open-access resource for pre-analyzed Arabidopsis network datasets. Overall, our work exemplifies the power of systems-level analyses and spatiotemporal transcriptomics to uncover hierarchical regulatory architectures in ETS. These findings not only redefine the functional scope of amino acid metabolism in bacterial pathogenesis but also position ANAC046 as a critical node in the susceptibility network, offering a potential target for engineering disease resistance.

## Results

### Comparative Coexpression Network Analysis Reveals ETS-Specific Rewiring of Amino Acid Metabolism in Arabidopsis

To determine ETS-dependent gene signatures and pathways, we performed comparative network analysis using WGCNA using high-resolution publicly available time-course transcriptome data from infection of Arabidopsis with DC3000, which induces ETS, and hrpA^−^, which elicits a robuts PTI response in the host (Lewis, Polanski et al. 2015, Mishra, Sun et al. 2017). The resulting comparative coexpression networks from these two interactions encompass 3,177 nodes and 15,724 edges (DC3000) and 1,881 nodes and 4,369 edges (hrpA^−^) **(Fig. 1, A and B; table S1)**. Similar to previously generated WGCNA-based coexpression networks in both animals and plants (He and Maslov 2016, Mishra, Sun et al. 2017, Naqvi, Zaidi et al. 2017, Mishra, Sun et al. 2018, Naqvi, Zaidi et al. 2019), both the DC3000 and hrpA^−^ coexpression networks exhibited scale-free characteristics when compared to respective random networks (r^2^=0.92) **(Fig. 1, C and D; table S1)**. Since deciphering network structural landscape and parameters of centrality measures may identify the most influential nodes within a network (Jeong, Mason et al. 2001, Brohee, Faust et al. 2008, Lü, Chen et al. 2016), we discovered that the average degree (nodes with high degree, *i.e.* number of links) and average betweenness (number of shortest paths that pass through a node) in the DC3000 network were significantly enriched with differentially expressed genes (DEGs) compared to hrpA^−^ **(Fig. 1, E and F; table S1)**. Given that the path length between pathogen effector targets and DEGs was previously shown to be shorter compared to comparable non-DEGs in an interactome (Mishra, Sun et al. 2017, Mishra, Kumar et al. 2019), clustering analysis of the coexpression network identified 17 and 16 different modules for DC3000 and hrpA^−^ networks, respectively **(Fig. 1, G and H and fig. S1 to S4; table S1)**. Moreover, we also demonstrated that the connectivity and cluster coefficient of modules in the DC3000 coexpression network (14.34 and 0.23, respectively) were significantly higher than in the hrpA^−^ coexpression network (6.42 and 0.22) **(fig. S1 to S4; table S1)**. Overall, the network topologies support a more complex regulatory network in cells of diseased plants compared to resistant plants. This can be explained by the regulatory effects of Type III effectors and other virulence molecules that target diverse plant processes.

**Fig. 1:**
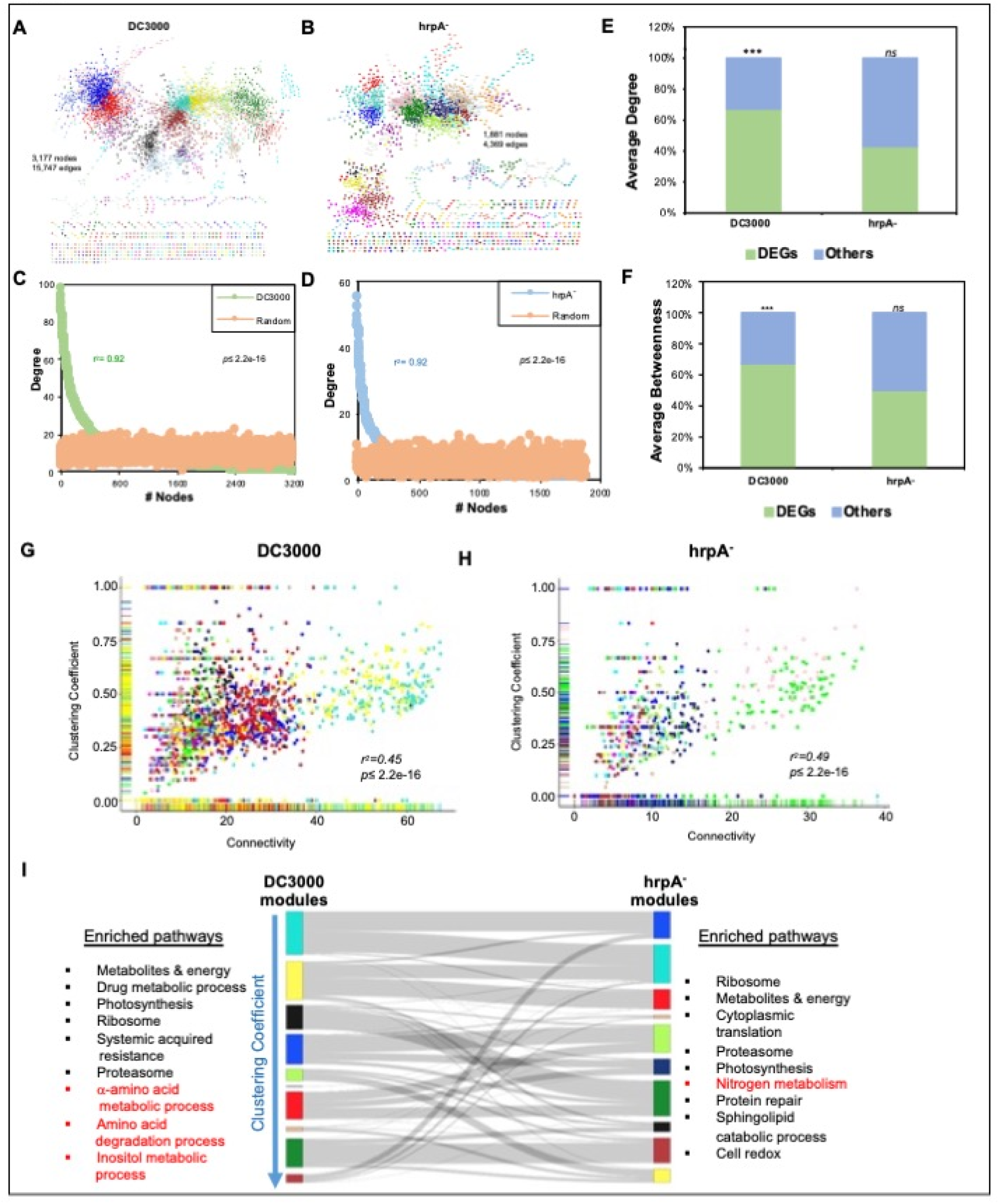
Gene coexpression network analysis revealed the high significance of differentially expressed genes (DEGs) and modules in effector-triggered susceptibility (ETS) vs. patterns-triggered immunity (PTI). **A.** *Arabidopsis*–*Pseudomonas syringae* pv. tomato DC3000 (hereafter DC3000) ETS gene coexpression network with 3,177 nodes and 15,724 edges represented by different module colors. B. *Arabidopsis* DC3000*hrpA*^−^ (hereafter hrpA^−^) PTI gene coexpression network with 1,881 nodes and 4,369 edges represented by different module colors. **C and D.** Node degree distribution of DC3000 and hrpA^−^ coexpression network follow power-law distribution (r^2^= 0.92) and their distribution is significantly different from their random network of the same size (*p*≤ 2.2e^-16^). **E.** Average degree of DEGs in DC3000 coexpression network is significantly higher (12.07) than other genes (non-DEGs; 6.0, *p*≤ 2.2e^-16^). DEGs in hrpA^−^ coexpression network have a lower (4.0) average degree than others (non-DEGs; 5.38). **F.** Average betweenness of DEGs in DC3000 coexpression network is significantly higher (0.0017) than for other genes (non-DEGs; 0.00087, *p*≤ 2.2e^-16^). DEGs in hrpA^−^ coexpression network have a higher (0.0011764) average betweenness than other genes (non-DEGs; 0.0011510) but the value is not statistically significant. **G.** The relationship between cluster coefficient and connectivity of DC3000 nodes illustrates a significant correlation (r^2^= 0.45, *p*≤ 2.2e^-16^). Turquoise, yellow, red, blue, and brown modules have significantly higher connectivity and cluster coefficient than any other modules. **H.** The relationship between cluster coefficient and connectivity of hrpA^−^ nodes illustrates a significant correlation (r^2^= 0.49, *p*≤ 2.2e-16). Green, pink, blue, and turquoise modules have higher connectivity and cluster coefficient than any other modules. **I.** Module-based similarity of DC3000 and hrpA^−^ common coexpressed genes along with enriched pathways. Modules are arranged based on their clustering coefficient in the DC3000 network.

Subsequently, we performed comparative coexpression analyses both at the qualitative (distinct nodes between two coexpression networks) and quantitative (common nodes between two coexpression networks that possess different clustering coefficients) levels. This allowed us to determine the coexpression landscape of ETS in DC3000-infected plants **(Fig. 1I; table S1)**. We then performed a comparative functional analysis to elucidate the common and distinct biological pathways pertinent to PTI and ETS. We observed several significantly enriched pathways shared between the DC3000 and hrpA^−^ coexpression networks, including photosynthesis, ribosome/translation, cell membrane, and cellular redox **(Fig. 1I and fig. S5)**. The unique enriched pathways in the PTI coexpression network were nitrogen metabolism and fructose metabolic pathways, whereas the significantly enriched pathways unique to the ETS coexpression network included the inositol metabolic process, α-amino acid metabolic process, and amino acid degradation process. These findings are consistent with physiological experiments demonstrating that plant metabolism is dramatically altered during ETS (O’Leary, Neale et al. 2016) and point to specific pathways that might support bacterial growth during infection.

We focused our investigation on amino acid-related pathways because these have not been previously implicated in ETS to bacteria **(Fig. 1I and fig. S5)**. We text-mined the *Arabidopsis* genome and extracted information for 165 amino acid-related genes (**table S2**). Within this group, 33 and nine unique genes were differentially regulated in response to DC3000 and hrpA^−^, respectively **(fig. S6; table S2)**. Among 55 differentially expressed genes that were shared between DC3000 and hrpA^−^, two groups could be discerned based on differences in transcriptional amplitude that indicted ETS-dependent regulation **(fig. S6; table S2)**. Moreover, we showed separate and distinct clusters of amino acid genes for ETS compared to PTI that were up- or down-regulated throughout pathogen infection. Collectively, these data indicate that DC3000 hijacks the amino acid-related pathways to establish disease susceptibility.

### Dynamic Regulatory Network Modeling Identifies ANAC046 as a Molecular Hub of ETS-Specific Amino Acid Gene Expression

Additional integral aspects of systems biology are analysis, simulation, and inference of temporal and cellular changes in gene regulatory networks as well as the construction of mechanistic models (Garbutt, Bangalore et al. 2014, Nobori, Velasquez et al. 2018, Mishra, Kumar et al. 2019). To better understand the dynamic transcriptional regulation of amino acid genes, we modeled ETS- and PTI-related gene expression datasets using the computational tool Scalable Models for the Analysis of Regulation from Time Series (SMARTS). This probabilistic graphical model integrates time-series expression with static protein-DNA interaction data to identify regulatory events and significant transcriptional regulators that exhibit bifurcated expression over time. Towards this, we first created a comprehensive database MID*ata* (*detailed below and Supplementary Materials*) of experimentally validated TF-target interactions (TFs: 1,151; Targets: 32,879; Interactions: 2,357,039). Modeling of ETS- and PTI-related transcriptomes featured major bifurcation events where the expression patterns of a cohort of genes differed from the remaining genes and revealed groups of TFs potentially governing specific sets of genes in a representative regulatory path **(fig. S7; table S2)**. Specifically, we modeled 78 and 33 different transcriptional paths that are controlled by 198 and 215 groups of TFs in ETS and PTI, respectively **(fig. S7; table S2)**. We identified a set of 82 distinct TFs that are specifically activated during the DC3000 infection but are absent from the PTI **(fig. S8a; table S2)**. These TFs could regulate plant pathways that promote pathogen virulence. We inferred underlying regulatory relationships between the 82 DC3000-induced TFs and amino acid-related genes, leading to 48 TFs predicted to directly control the expression of 123 amino acid-related genes during ETS **(Fig. 2A and fig. S8b; table S2)**. Among these, we predicted the TF ANAC046 (Arabidopsis NAC [NAM (no apical meristem), ATAF1/2, CUC2 (cup-shaped cotyledons 2)] 046) as a candidate master regulator that might influence the amino acid metabolism and transport during pathogen infection. ANAC046 was previously predicted as a major regulatory hub (i.e. highly connected node) in networks regulating development and senescence(Mishra, Sun et al. 2018). To comprehensively examine the interactions of ANAC046 within coexpression and gene regulatory networks (GRNs), we queried the curated MID*ata* database using its locus ID (AT3G04060) for both coexpression and TF–target relationships. This analysis revealed that ANAC046 is coexpressed with a total of 941 genes, including 32 in the DC3000 network, 279 in the developmental biology network (senescence), and 310 across 16 distinct hormone-responsive networks **(Fig. 2A; table S3)**. Moreover, based on TF–target datasets including Cistrome, ANAC046 is predicted to directly regulate up to 10,572 genes **(Fig. 2A; table S3)**, highlighting its broad regulatory potential. Interestingly, ANAC046 was not present in the PTI coexpression network, suggesting that it may function as a regulatory hub specifically in ETS rather than PTI **(Fig. 1A, B; table S3)**.

**Fig. 2.**
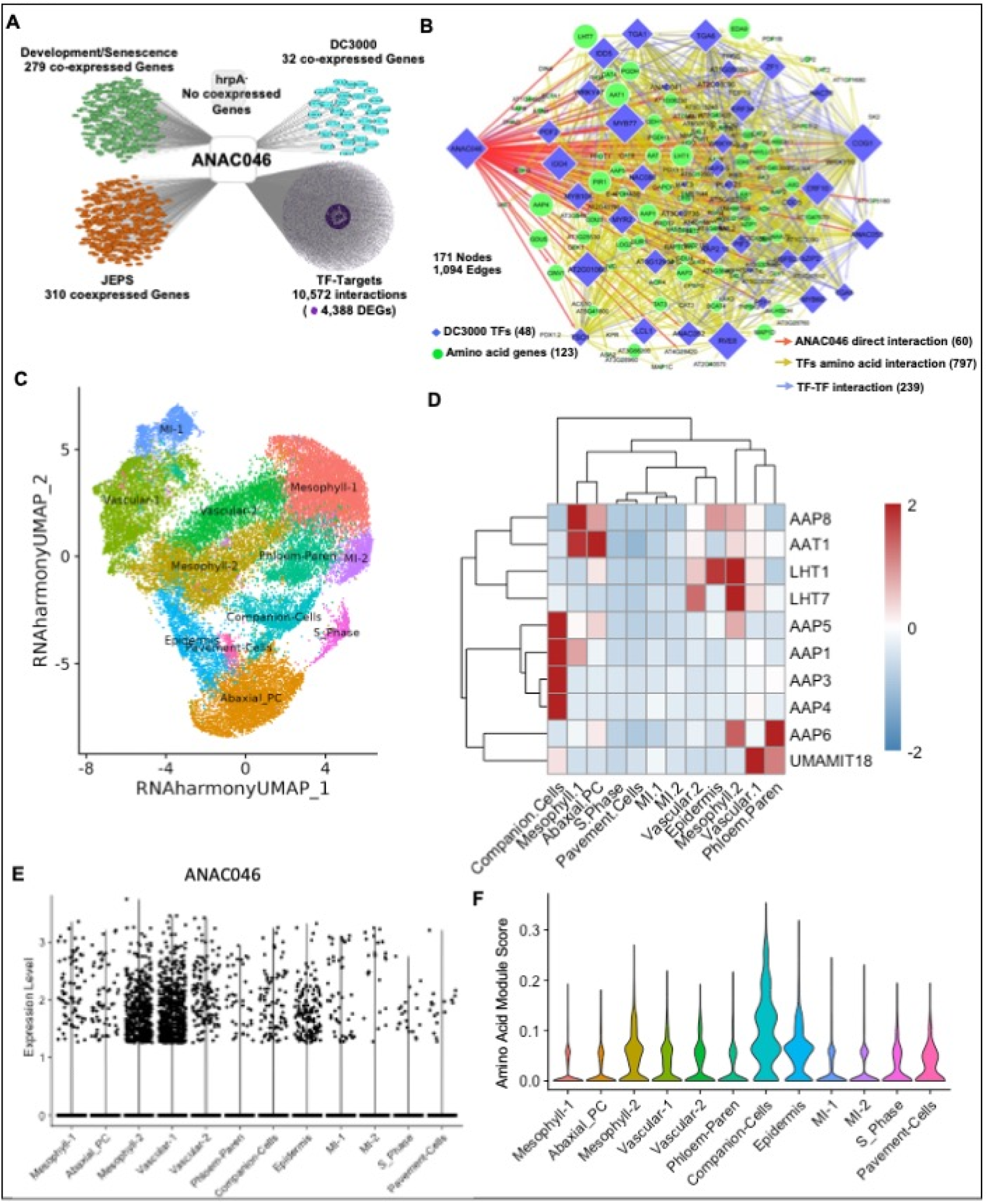
ANAC046 regulates the expression of amino acid genes during *Pst* DC3000 infection. **A.** MIData database extracted interactions including gene coexpression networks and TF-Target interactions of ANAC046. **B.** ANAC046 directly/indirectly regulates amino acid transporter/metabolism genes (123 green circles) and transcription factors (TFs) (48 blue squares including ANAC046), which regulate amino acid transporter genes. ANAC046 direct regulatory interactions are represented by light red arrows while other TF- amino acid transporter gene interactions are represented by yellow arrows. TF-TF interactions are marked by light blue arrows. The size of the node represents the degree of a node in the DC3000 coexpression network (Fig. 1A). All possible TF-Target relationships (10,445 interactions) of ANAC046 were extracted from the <u>MID*ata* database</u>. **C.** The single cell RNA-Seq (scRNA-Seq) analysis UMAP representation of annotated cell types for publically available dataset GSE226826. **D**. The gene expression heatmap of NAC046 regulated amino-acid transporters in different cell types in scRNA-Seq. The red color indicates high expression, while the blue color represents low expression. **E**. The expression of ANAC046 in different cell types of leaf scRNA-Seq data. **F.** The amino acid module scoring in different cell types form two independent single cell datasets. UCell Module scoring function with genes from Figure 2D was used to perform the scoring analysis.

Next, we extracted a targeted GRN subnetwork centered on 48 transcription factors (TFs)—including ANAC046 and 47 additional TFs—and 123 amino acid-related genes. This analysis revealed 837 TF–gene interactions, of which 40 interactions are directly mediated by ANAC046 and 797 by the remaining TFs **(fig. S8a; table S3)**. Subsequently, we mapped potential regulatory interactions among the 48 TFs themselves to identify TF–TF interactions. This analysis revealed 239 such connections, including 20 direct interactions between ANAC046 and other TFs. Integrating these layers, we constructed a comprehensive ANAC046 amino acid GRN comprising 171 nodes (123 amino acid genes and 48 TFs) and 1,094 interactions: 60 ANAC046-directed edges (40 to amino acid genes and 20 to TFs), 797 edges from other TFs to amino acid genes, and 239 TF–TF edges **(Fig. 2B; table S3)**. These findings further support ANAC046 as one of the major regulatory hub, prominently modulating susceptibility-associated transcriptional programs during ETS.

To investigate the cellular expression patterns of amino acid-related genes and their potential regulation by ANAC046, we performed scRNA-Seq analysis using recently published, time-resolved datasets from Arabidopsis leaves infected with DC3000 (GSE226826)(Nobori, Monell et al. 2025). The scRNA-Seq analysis revealed 12 cell clusters classified as two mesophyll, two vascular, two MI, two pavements cells, pholem parenchema, epidermis, and s-phase cell populations **(Fig. 2C, fig. S9a-c)**. Interestingly, we identified that most of the amino acid transporters and significantly expressed in companion cells and mesophyll cells **(Fig. 2D, fig S9d)**. Subsequently, we examined the expression patterns of amino acid transporters—specifically members of the AAP, UMAMIT, LHT, and AAT families—with a focus on those directly targeted by ANAC046. This subset includes five AAP transporters (*AAP1*, *AAP3*, *AAP4*, *AAP5*, and *AAP6*), two LHT transporters (*LHT1* and *LHT7*), and two additional transporters, USUALLY MULTIPLE ACIDS MOVE IN AND OUT TRANSPORTERS 18 (*UMAMIT18)* and AMINO ACID TRANSPORTER 1 (*AAT1)*. Intriguingly, we found that the ANAC046 direct target AAP transporters are primarily expressed in companion cells, whereas *AAP8*—an AAP family member not directly targeted by ANAC046—is predominantly expressed in mesophyll 1 and abaxial pavement cells. In contrast, *AAP6*, *LHT1*, and *LHT7* show highest expression in mesophyll 2 cells. Additionally, *UMAMIT18* is mainly expressed in vasculature 1, while *AAT1* is primarily expressed in mesophyll 1 cells. We also found that ANAC046 is expressed across diverse cell types, consistent with its role as a regulatory hub. Its expression is particularly elevated in mesophyll and vascular cells, as observed in dataset GSE226826(Nobori, Monell et al. 2025) **(Fig. 2E)**. To further validate the impact of amino acid transporters in single cell dataset, we calculated the amino acid module scoring by UCell. This analysis validated the high scoring for amino acid in companion cells in GSE226826(Nobori, Monell et al. 2025) **(Fig. 2F)** and in another Arabidopsis leaf scRNA-Seq dataset GSE213625(Zhu, Lolle et al. 2023) **(fig S9e)**. Integrating literature-based experimental evidence, transcriptional simulations, and scRNA-seq expression patterns, we conclude that ANAC046 directly regulates members of multiple amino acid transporter families, along with key enzymatic genes. Based on these findings, we further hypothesize that DC3000 targets ANAC046 to promote full virulence.

To test this hypothesis, we first examined the temporal dynamics and amplitude of target gene mRNAs in response to DC3000 and hrcC^−^ in Col-0 and *ANAC046-SRDX* plants. These experiments confirmed ANAC046 transcriptional dependency of selected 10 direct target genes by demonstrating a significantly increased or decreased transcript abundance in the *ANAC046-SRDX* mutant at one or more time points following treatment with both bacterial strains under study **(Fig. 3; table S4)**. Moreover, ANAC046-dependent genes displayed differential transcript accumulations upon infection with DC3000 compared to hrcC^−^ **(Fig. 3; table S4)**. Although *AAP8* and *LHT4* are not direct targets of ANAC046, they were included in our experiments because other members of the AAP and LHT families are regulated by ANAC046. Notably, these two indirect targets exhibited distinct expression patterns compared to their respective family members that are directly regulated by ANAC046. Notably, amino acid metabolism and transporter genes have previously been implicated in plant–pathogen interactions(Yao, Sui et al. 2025). Here, we compiled a list of pathology-related phenotypes for these genes (**table S4**). These experiments validate the predicted role of ANAC046 as a master regulator of amino acid-related genes during ETS (**table S4**).

**Fig. 3:**
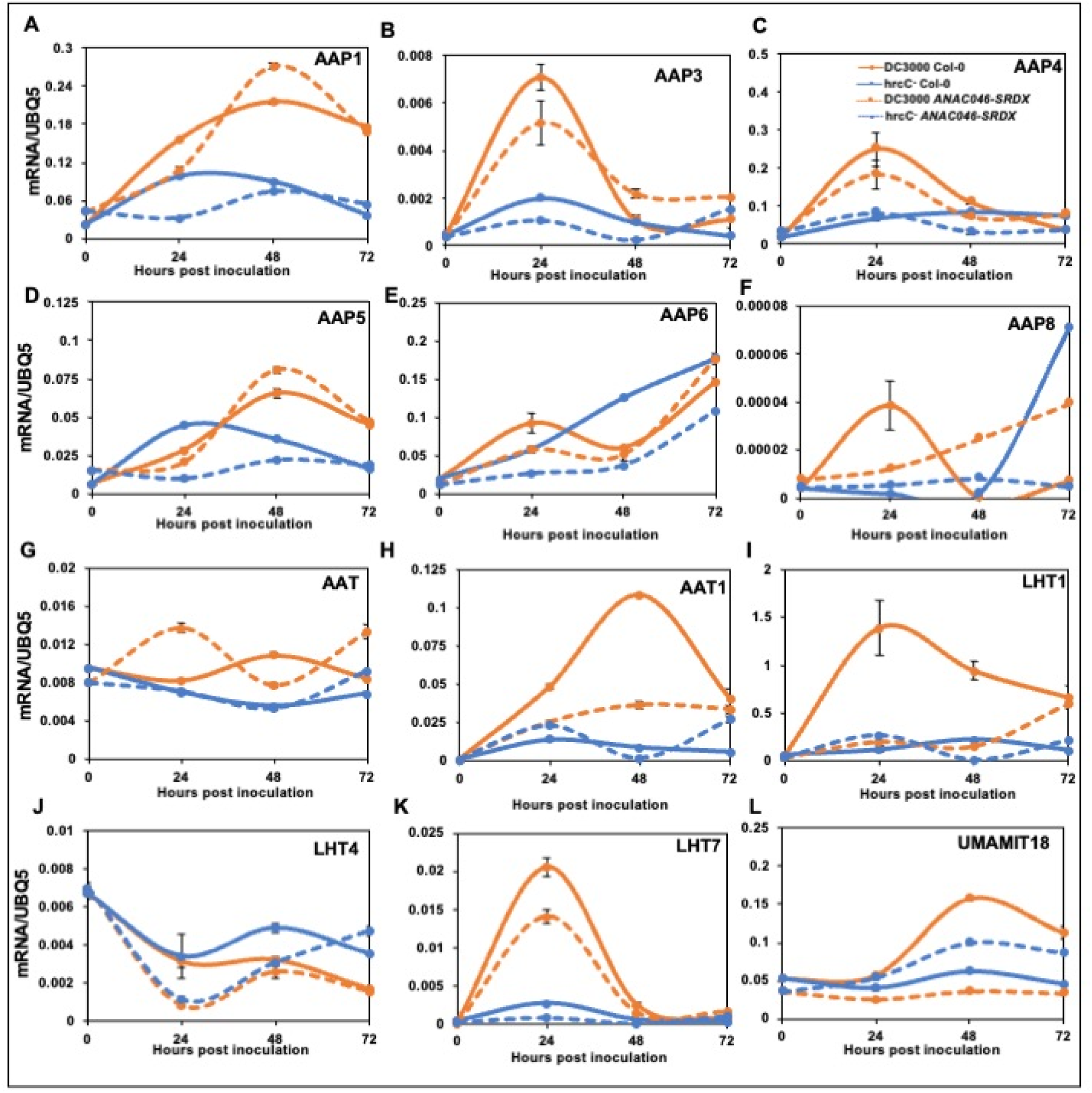
Kinetics of gene expression of *ANAC046*-related amino acid transporters in Col-0 and *ANAC046-SRDX*. Quantitative reverse transcription-polymerase chain reaction (RT-qPCR) was performed in Col-0 and *ANAC046-SRDX* genotypes upon treatment with DC3000 and hrcC^−^ at 0h, 24h, 48h, and 72h. Gene expression was assessed using reference gene *UBQ5* in (**A**) *AAP1*, (**B**) *AAP3*, (**C**) *AAP4*, (**D**) *AAP5*, (**E**) *AAP6*, (**F**) *AAP8*, (**G**) *AAT*, (**H**) *AAT1*, (**I**) *LHT1*, (**J**) *LHT4*, (**K**) *LHT7*, and (**L**) *UMAMIT18*. The graph represents the mean with standard errors of two technical replicates.

To further test the hypothesis that DC3000 targets ANAC046 to promote its virulence, we monitored ANAC046 transcript accumulation on a time course following infection of *Arabidopsis* with DC3000 or the non-pathogenic DC3000 mutant hrcC^−^. Like HrpA^−^, hrcC^−^ is defective in type III secretion and cannot deliver effector molecules to the host cells. We showed that *ANAC046* mRNA is steadily induced in response to DC3000 over 72 hours following infection **(Fig. 4A)**. Contrastingly, *ANAC046* is not induced by the hrcC^−^ strain, consistent with a function in ETS but not PTI. To substantiate the role of ANAC046 in ETS, we employed *ANAC046-SRDX* transgenic plants, which express a chimeric repressor containing the SRDX domain and exhibit a loss-of-function phenotype of ANAC046(Mishra, Sun et al. 2018). *ANAC046-SRDX* displayed an enhanced disease resistance phenotype to DC3000 but not hrcC^−^ **(Fig. 4B)**, providing genetic evidence that ANAC046 is necessary for full susceptibility to DC3000 but is irrelevant to the interaction with hrcC^−^. Consistent with these disease resistance phenotypes, we also demonstrated an increased mRNA accumulation of two well-characterized salicylic acid-responsive molecular markers, *pathogenesis-related 1* (*PR1*) and *pathogenesis-related 5* (*PR5*), in the *ANAC046-SRDX* transgenics compared to Col-0 in response to DC3000 **(Fig. 4C and D; table S4)**. In contrast, the DC3000-dependent transcript accumulation of jasmonic acid pathway markers *oxophytodienoate reductase 3* (*OPR3*) and *jasmonate-ZIM-domain 5* (*JAZ5*) was dampened in the *ANAC046-SRDX* transgenics **(Fig. 4E and F; table S4)**. While we observed a significant reduction of *flg22-induced receptor-like kinase 1* (*FRK1*) mRNA in an ANAC046-dependent manner **(Fig. 4G; table S4)**, the plants lacking functional ANAC046 are not compromised in flg22-induced PTI **(Fig. 4H; table S4)**. Collectively, these data indicate that ANAC046 is not necessary for PTI and that DC3000 requires ANAC046 to establish efficacious ETS. Overall, these experiments provide evidence that ANAC046 functions as a master regulator of amino acid-related genes in plant defense.

**Fig. 4:**
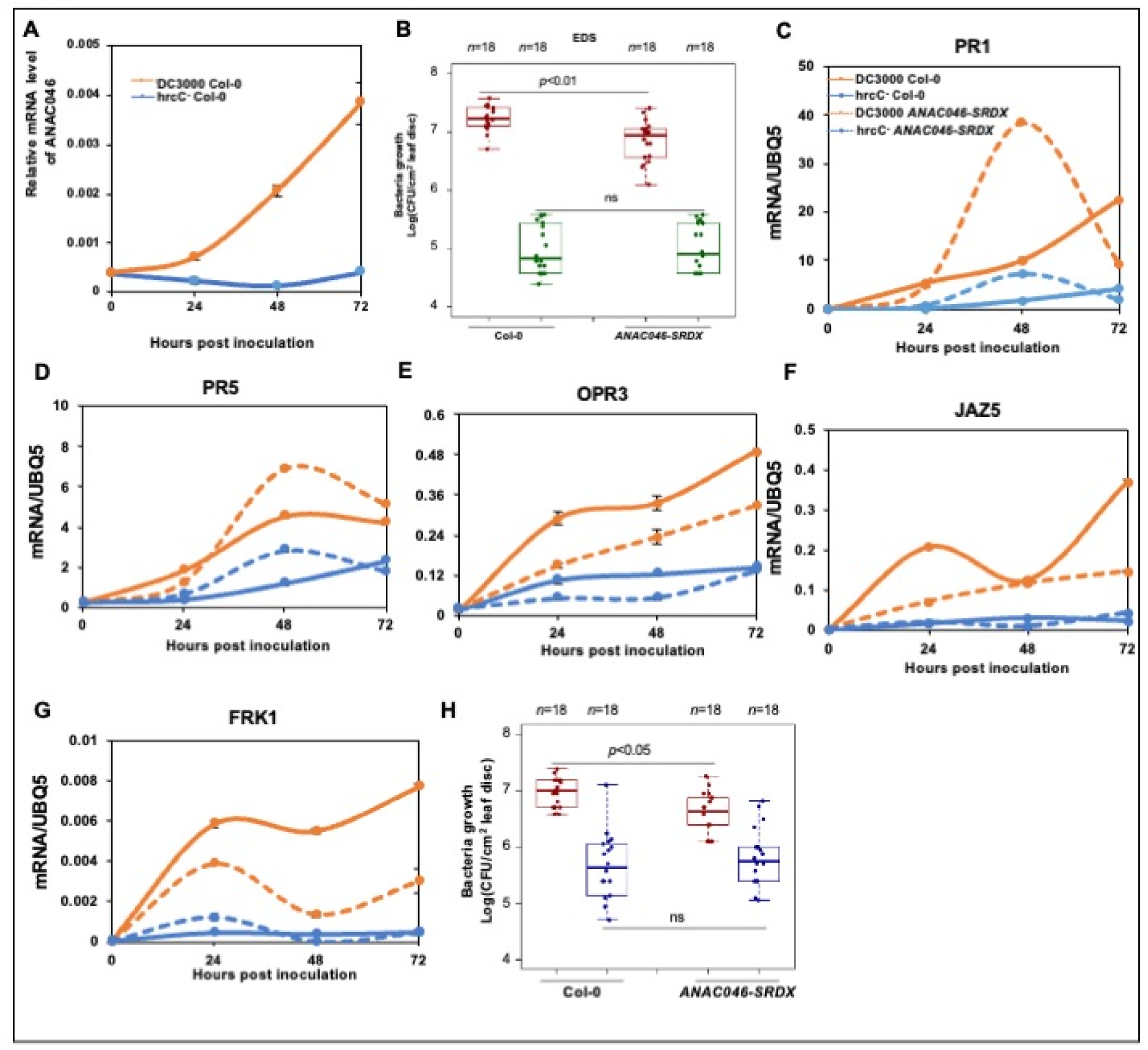
Kinetics of gene expression of *ANAC046* and defense-responsive markers under different pathogen treatments. **A.** Kinetics of gene expression of *ANAC046* in Col-0 in response to infection with DC3000 and hrcC^−^at 0hr, 24hrs, 48hrs and 72hrs post pathogen inoculation. **B.** Enhanced Disease Resistance (EDR) of *ANAC046-SRDX*. Bacterial growth of DC3000 (red bars) and hrcC^−^ (a type III secretion system mutant strain of DC3000, green bars) were quantified 3 days after syringe inoculation (OD_600nm_ = 0.0002) on *ANAC046-SRDX*. Col-0 plants were used as controls. Dots in the box and whisker plots represent individual data points. “n” corresponds to the number of leaf samples, and each sample contains three biological independent leaf discs. **C-F.** Kinetics of gene expression of *ANAC046*-related markers in Col-0 and *ANAC046-SRDX*. Quantitative reverse transcription-polymerase chain reaction (RT-qPCR) was performed in Col-0 and *ANAC046-SRDX* genotypes upon treatment with DC3000 and hrcC^−^ at 0h, 24h, 48h, and 72h. Gene expression was assessed using reference gene *UBQ5* in (**C**) *PR1*, (**D**) *PR5*, (**E**) *OPR3*, (**F**) *JAZ5*, (**G**) *FRK1*. The graph represents the mean with standard errors of two technical replicates. **H.** Patterns-triggered immunity (PTI) of *ANAC046-SRDX*. Red and blue whisker-box plots represent plants infected with DC3000 and flg22 pretreatment followed by DC3000, respectively. Bacterial growth was quantified 3 days after needleless syringe inoculation (OD_600nm_ = 0.002). The genotypes Col-0 and *ANAC046-SRDX* are indicated. Each dot in the box and whisker plot represents an individual data point. “n” represents the number of leaf samples, and each sample contains three independent leaf discs. For **B** and **H** One-way analysis of variance (ANOVA) was performed to estimate statistical significance for bacteria growth. n.s. stands for not significant. \**p* < 0.05, \*\**p* < 0.01, and \*\*\**p* < 0.001.

### Inner Network Layers of the Plant Interactome Enriched in Amino Acid Metabolism

Another important tenet of systems biology is the integration of multi-omics data including transcriptome and interactome, which may provide valuable insights into the complexity of plant-pathogen interactions (Subramanian, Verma et al. 2020). Towards this, we first generated the largest to-date <u>p</u>rotein-<u>p</u>rotein interactions <u>e</u>xperimental (PPIE) network by combining all the existing large- scale interactomes as well as literature-curated interactions (LCI) **(Fig. 5A-E; table S5)**. The PPIE network encompasses 10,107 nodes and 40,027 edges as well as 869 hubs (highly connected nodes) and exhibits a scale-free network property **(Fig. 5F; table S5)**. Since network centrality has been demonstrated to hold high predictive power for identifying significant molecular players (Ahmed, Howton et al. 2018, Mishra, Kumar et al. 2019, Subramanian, Verma et al. 2020), we showed a positive correlation between betweenness and degree (r^2^=0.54) as well as betweenness and stress (r^2^=0.73) **(Fig. 5G and H; table S5)**. Previously, the weighted *k*-shell decomposition method was shown to significantly increase the predictive power of host-pathogen targets to 40% in the inner layers of two independent interactomes (Ahmed, Howton et al. 2018). Thus, we subjected the PPIE network to weighted *k*-shell decomposition analysis and identified 2,539 inner and 7,568 outer layer nodes **(Fig. 5J; table S5)**. Consistent with previous network topology analyses of other interactomes (Ahmed, Howton et al. 2018), we found that the degree and distance from the core displayed power-law correlation **(Fig. 5I; table S5)**. Consequently, the average degree, clustering coefficients, and stress in the nodes residing in the inner layers of the PPIE network are significantly heightened compared to that of the nodes in the peripheral layers **(Fig. 5 K-M; table S5)**. To explore the functional role of inner layers, we performed gene ontology (GO) and KEGG pathway analyses and showed that the inner node proteins were enriched in “amino acid biosynthesis” as well as “hormone signal transduction”, “carbon metabolism” and “protein processing in the Endoplasmic Reticulum (ER)” GO categories **(Fig. 5J)**. Given that pathogen effector targets are preferentially located in the inner layers of an interactome exhibiting increased degree and betweenness, and the inner layers’ nodes of the PPIE network display similar features, this model underscores the biological importance of amino acid metabolism genes in plant-pathogen interactions.

**Fig. 5:**
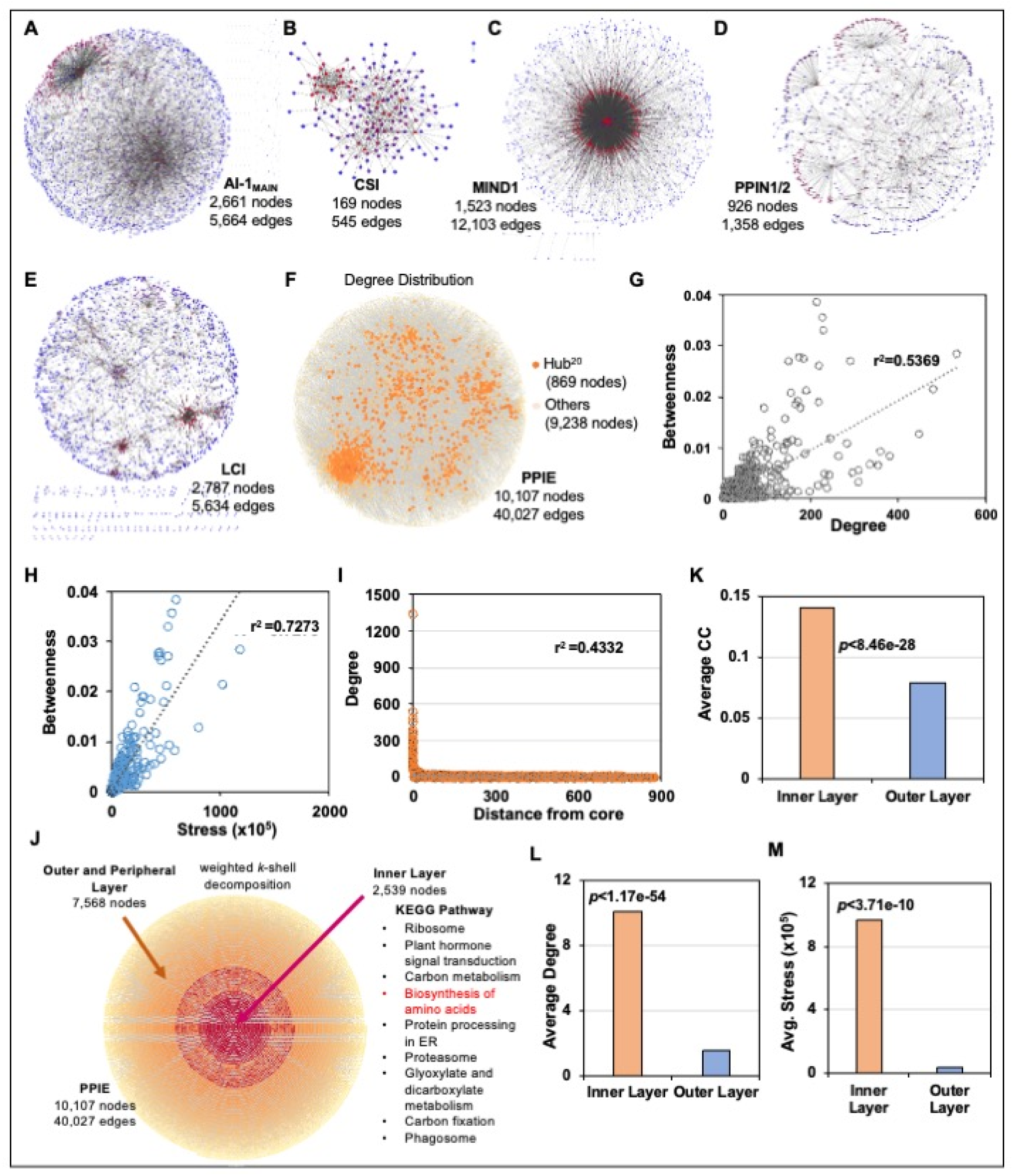
Network analyses of diverse *Arabidopsis* interactomes. **A.** *Arabidopsis* interactome 1 (AI-1MAIN) with 2,661 nodes and 5,664 edges; **B.** *Arabidopsis* cell surface interactome (CSI) with 169 nodes and 545 edges; **C.** Membrane-Linked Interactome of *Arabidopsis* (MIND1) with 1,523 nodes and 12,103 edges; **D.** *Arabidopsis* Plant-Pathogen Immune Network (PPIN1) with 961 nodes and 1,358 edges; **E.** *Arabidopsis* literature curated interactions (LCI) with 2,787 nodes and 5,634 edges; **F.** Combined *Arabidopsis* protein-protein experimental interaction (PPIE) network with 10,107 nodes and 40,027 edges (yellow-red nodes: 869 Hub^20^ proteins, light yellow nodes: 9,238 other proteins); **G.** Distribution of degree and betweenness centrality in PPIE network (r^2^= 0.54); **H.** Distribution of stress centrality and betweenness centrality in PPIE network (r^2^= 0.7273); **I.** Distribution of degree and distance from the core of the network in PPIE network (r^2^= 0.4332); **J.** Weighted *k*-shell decomposition of combined with 10,107 nodes and 40,027 edges in PPIE network (Red nodes: 2,539 Inner layer proteins, Blue nodes: 7,568 Outer/peripheral layer proteins). The enriched Kyoto Encyclopedia of Genes and Genomes (KEGG) pathways of inner layer proteins are listed. **K-M.** Average cluster coefficient (CC), average degree, and average strees of inner layer proteins is significantly higher than outer layer proteins (*p* ≤ 8.46e^-28^, 1.17e^-54^, 3.71e^-10^, respectively).

### Multi-Omics Integration and Network-Driven Functional Discovery Links Amino Acid Metabolism and Nitrogen Signaling to Plant Defense

To comprehensively understand the relationship between amino acid-related genes and ETS or PTI, we integrated the PPIE network with a high-resolution time-course transcriptome corresponding to DC3000 and hrpA^−^ with 5,141 nodes and 12,813 edges, and the differential gene expression in companion cells during pathogen infection (DC3000 vs. mock) from a single cell RNA-Seq study **(Fig. 6A; fig. S10a; table S5).** We observed that the hrpA^−^-derived network (558 nodes and 208 edges) was smaller than the DC3000 network (1,762 nodes and 1,501 edges), similar to the comparative coexpression networks described earlier. A substantial component of the PPIE was regulated in response to both DC3000 and hrpA^−^ and encompassed 2,821 nodes and 3,732 edges **(Fig. 6A; fig. S10a; table S5).** We also identified a significant overlap in the companion cells DEGs among inner layers of PPIE and the 1^st^ degree and 2^nd^ degree interactors of amino acid genes **(Fig. 6A, fig. S10b)**. Furthermore, we determined that 142 direct interacting partners of proteins corresponding to amino acid-related genes (1^st^ neighbors) were enriched in GO categories such as “plant-pathogen interaction”, “extracellular stimuli”, chemical homeostasis, “ion transport” and “defense response” **(fig. S10c)**. Intriguingly, these 1^st^ neighbors of amino acid-related genes are enriched with hubs compared to 2^nd^ neighbors in the PPIE network (hypergeometric *p* < 1.47e-48), further substantiating the critical importance of amino acid-related genes within a network as well as the significance of these genes in plant-microbe interactions **(Fig. 6A; table S5).** Taken together, our comparative analyses of two plant defense-centered coexpression networks and topological analyses of an integrative transcriptome-interactome PPIE network independently point to the potential functions of amino acid-related genes in plant immune responses. Accordingly, a subset of genes involved in amino acid transport has been implicated in plant disease resistance and susceptibility to a wide spectrum of pathogens including bacteria, fungi, oomycetes, and nematodes, providing a proof-of-concept to our systems biology-aided discovery **(table S4)**.

**Fig. 6:**
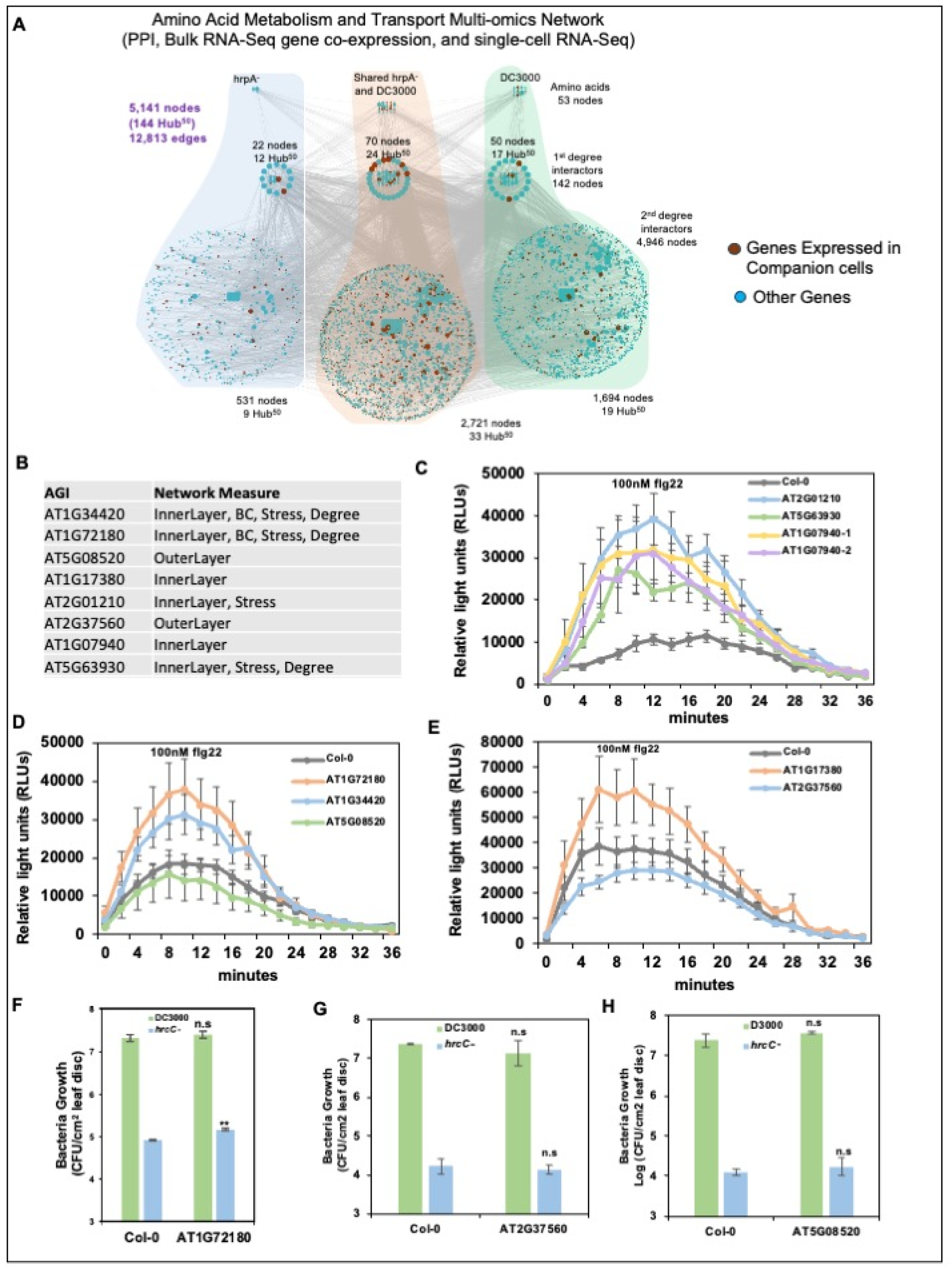
Multi-omics data integration identifies novel plant defense players in *Arabidopsis*. **A.** 53 differentially expressed amino acid genes in DC3000 and hrpA^−^ networks interact with 5,088 additional expressed proteins corresponding to 12,813 edges. We identified 142 1^st^ degree and 4,946, 2^nd^ degree interactors of amino acid proteins. Of these 142 1^st^ degree interactions, 22, 50, and 70 proteins are differentially expressed in hrpA^−^, DC3000 and shared between both conditions, respectively, while 531, 2,721, and 1,694 proteins are 2^nd^ degree interactors. Additionally, 12, 24, and 17 proteins of 1^st^ degree interactors are Hub^50^ in hrpA^−^, DC3000 and shared between both conditions, respectively. Next, 9, 33, and 19 proteins of 2^nd^ degree interactors are Hub^50^ in hrpA^−^, DC3000 and shared between both conditions, respectively. Finally, we mapped the companion cells differential expressed genes during pathogen infction (DC3000) from the single cell RNA-Seq study; **B.** Selected hub genes for experimental analysis; **C-E.** Reactive oxygen species (ROS) burst was measured using a luminol-based assay immediately after treatment of plants with 100mM flg22 for 36 minutes. Four-week-old *Arabidopsis* leaves were used to punch leaf discs. ROS profile of Col-0, **C.** CS860541 (AT2G01210), SAIL_900_B07 (AT5G63930), SALK_049659C (AT1G07940-1) and SAIL_620_H05 (AT1G07940-2); **D**. SALK_081193C (AT1G72180), SALK_109214C (AT1G34420) and SALK_205581C (AT5G08520); **E.** SALK_053776 (AT1G17380), SALK_086357C (AT2G37560); was shown as mean ± SE from 8 leaf discs. **F-H.** Bacterial growth of DC3000 (green bars) and *hrcC^−^*(blue bars) were quantified 72 hours after needleless syringe inoculation with OD_600nm_= 0.02 on **F.** SALK_025328C (AT2G36720), SALK_081193C (AT1G72180), **G.** SALK_086357C (AT2G37560), **H.** SALK_205581C (AT5G08520). The results shown are mean ± SE. n=6, \**p*<0.05, \*\**p*<0.01, \*\*\**p*<0.001, n.s. stands for non-significant according to two-tailed Student’s t-test.

Since network architectural and topological analyses can identify significant players in a biological system, particularly plant-pathogen interactions (**Fig. 6B**) (Crua Asensio, Munoz Giner et al. 2017, de Groot and Torrent Burgas 2020), we sought to further unravel the plant immune functions for seven additional nodes that exhibit a high degree, stress and betweenness centrality, and reside within the inner layers of the PPIE network (**table S5**). In parallel, we included two additional genes, *AT5G08520* and *AT2G37560*, as comparative controls. These genes are positioned in the peripheral (outer) layers of the network and display relatively low centrality metrics. We acquired homozygous loss-of-function mutants for these genes and subjected them to a flg22-induced reactive oxygen species (ROS) burst assay. All seven mutant lines corresponding to inner-layer nodes with elevated centrality metrics exhibited a significantly enhanced ROS burst when compared to Col-0. In contrast, the two control mutants (*AT5G08520* and *AT2G37560*), which are localized in the outer layers of the PPIE network and lacked high centrality characteristics, did not show any significant difference in ROS production relative to Col-0 **(Fig. 6C - E; table S4)**. Subsequently, we focused on *AT1G72180* (aka *CEPR2*; C-terminally encoded peptide receptor 2), which is a leucine-rich repeat receptor kinase that functions as a receptor for a novel peptide phytohormone, CEP1. *CEPR2* directly interacts with the ABA transporter NRT1.2/NPF4.6 and mediates nitrate uptake signaling (Zhang, Yu et al. 2021). We tested the mutants for these genes for their responses to DC3000 and hrcC^−^, and demonstrated that loss-of-function mutant *cepr2* supported increased bacterial load to hrcC^−^ only **(Fig. 6F; table S4)**. Finally, two mutants corresponding to control genes *AT5G08520* and *AT2G37560* did not show any significant change in the ROS production or bacterial susceptibility **(Fig. 6C - E**; **Fig. 6G and H; table S4**). Taken together, our network topology analyses discovered seven previously uncharacterized Arabidopsis nodes involved in the plant immune system, one of which is linked to nitrogen.

### Creation of MID*ata*: A Comprehensive Multi-Omics Integration Platform for *Arabidopsis thaliana*

The success of our large-scale integrative multi-omics analyses motivated us to construct a global platform **M**ulti-omics **I**ntegrated **D**atabase of ***A****rabidopsis **t**halian**a*** (MID*ata*) for use by the plant research community. We curated and analyzed data acquired from diverse *Arabidopsis* -omics experiments including gene coexpression networks, TFs and target genes networks, PPI, and phonemics networks. The current version of MID*ata* 1.0.0 contains over 22,000 *Arabidopsis* coexpressed genes, over 3,500,000 TFs and target gene pairs, >21,000 PPI data with nodes, and more than 6,000 genes with phenotype key terms brought together through rigorous curation **(fig. S11 a-b)**.

The MID*ata* database can be accessed at https://midata.cas.uab.edu/. The user-friendly interface also provides information on network centrality analyses including degree, betweenness, and closeness as well as a downloadable link with all of the -omics and network analyses information for a queried *Arabidopsis* Genome Initiative identifier (AGI ID). MID*ata* provides a global view of how different - omics interactions relate to a phonotype and allows the understanding of information flows among various layers of -omics-network. We anticipate that MID*ata* will fill an important niche and accelerate the adoption of systems biology approaches in the plant research community. We also generated a Cytoscape plugin app to perform weighted *k*-shell decomposition analysis (http://apps.cytoscape.org/apps/wkshelldecomposition**, fig. S11c)**.

## Discussion

Plants rely on a multilayered immune system composed of PTI and ETI. Although significant advances have been made in cataloging transcriptional responses triggered by these immune layers, a comprehensive understanding of how pathogens manipulate host regulatory networks at a systems level has remained limited. Our study addresses this gap by offering a global view of how ETS reprograms host cellular functions, with a specific emphasis on amino acid metabolism—a metabolic pathway that has been relatively underexplored in the context of plant-pathogen interactions. Using integrative multi-omics analyses, we identified ANAC046 as a master regulator of amino acid-related genes. Our network science–driven analyses further revealed that amino acid–related genes are significantly enriched in network hubs and led to the discovery of previously uncharacterized genes involved in plant defense.

Our integrative multi-omics analyses, encompassing coexpression networks, regulatory modeling, and single-cell transcriptomics, have unveiled significant transcriptional and regulatory reprogramming in Arabidopsis during ETS, particularly affecting amino acid metabolism and nutrient dynamics **(Fig. 1)**. Amino acids play multifaceted roles in plant-pathogen interactions, serving as both nutrients and signaling molecules. Studies using transcriptomics and metabolite profiling in Arabidopsis have shown that infection by both virulent and avirulent strains of DC3000 as well as PTI elicitors triggers significant changes in amino acid-related gene expression and metabolite levels(Rojas, Senthil-Kumar et al. 2014). For instance, avirulent strains increase levels of tryptophan, tyrosine, lysine, valine, and leucine while decreasing glutamate, whereas virulent strains lead to higher levels of isoleucine, threonine, alanine, phenylalanine, tyrosine, and glutamine. Aspartate levels decrease in both cases(Rojas, Senthil-Kumar et al. 2014). These findings suggest that particular amino acids may be involved in plant defense responses, although further studies are needed to confirm their specific roles. Complementing this metabolic framework, amino acid transporters—such as AAPs, UMAMITs, LHTs, AATs families—play crucial roles in modulating plant-pathogen interactions by regulating the transmembrane transport of amino acids(Sonawala, Dinkeloo et al. 2018, Yao, Sui et al. 2025). These transporters can either enhance plant defense by controlling nutrient allocation or, alternatively, be exploited by pathogens to access amino acids needed for their growth and virulence. Our study revealed that specific AAPs, such as *AAP1*, *AAP3*, *AAP4*, and *AAP5*, are significantly up-regulated during ETS, suggesting that pathogens may exploit these transporters to facilitate infection by manipulating host nutrient fluxes **(Fig. 2 and Fig. 3)**. Interestingly, *AAP6* and *AAP8* show early induction during ETS, but their expression becomes prominent during the later stages of PTI, indicating a nuanced role in the transition between susceptibility and immune activation. This expression pattern aligns with earlier findings that AAPs can act as susceptibility genes (**table S4**), particularly in the context of nematode infections(Elashry, Okumoto et al. 2013, Yao, Sui et al. 2025). Several AAPs have been shown to be induced upon nematode infection, potentially facilitating the transfer of amino acids to nematode feeding sites, thereby supporting pathogen growth(Elashry, Okumoto et al. 2013, Dutta, Rupinikrishna et al. 2024). Notably, all Arabidopsis AAPs except *AAP5* and *AAP7* are up-regulated in response to nematode infection, reinforcing the idea that AAP5 may play a distinct role in immune modulation(Yao, Sui et al. 2025). Functional analyses of *aap1*, *aap2*, *aap3*, *aap6*, and *aap8* mutants reveal significantly reduced nematode development and reproduction, further corroborating their roles as facilitators of pathogen success(Elashry, Okumoto et al. 2013, Yao, Sui et al. 2025).

Complementing this, the UMAMIT family—primarily functioning as amino acid exporters—also participates in defense and susceptibility pathways. In our study, *UMAMIT18* is upregulated during ETS **(Fig. 2 and Fig. 3)**, suggesting a potential link between pathogen manipulation of host amino acid transport and disease progression. This upregulation was not observed upon activation of PTI, indicating that *UMAMIT18* is not broadly responsive to immune signals but may be specifically co-opted or dysregulated during effector-mediated processes. Previous studies have demonstrated that members of this family, such as *UMAMIT14* and *UMAMIT18*, contribute to radial amino acid transport in roots, supporting amino acid secretion into the rhizosphere (Besnard, Pratelli et al. 2016). These secreted amino acids can influence root–microbe interactions, a relationship further emphasized by the role of *LHT1*, an amino acid importer, in modulating the composition of root exudates to affect the growth and metabolic activity of rhizosphere microbes such as *Pseudomonas simiae* (Agorsor, Kagel et al. 2023). *LHT1* has also been characterized as a key importer of amino acids from the leaf apoplast, crucial for maintaining intracellular amino acid pools(Yao, Sui et al. 2025). Intriguingly, *LHT1* expression is upregulated during ETS, a phase typically associated with pathogen proliferation and suppression of plant immunity. This observation aligns with previous findings that knockout of *LHT1* enhances resistance to a broad spectrum of pathogens (Rogan, Pang et al. 2024). The resistance phenotype in *lht1* mutants has been linked to glutamine deficiency and alterations in cellular redox status, which in turn activate salicylic acid (SA)-dependent defense pathways. Adding further complexity, *LHT7*—a transporter closely related to *LHT1*—is also upregulated in ETS, suggesting a coordinated manipulation of amino acid transport during pathogen infection **(Fig. 2** and **Fig. 3)**. In contrast, *LHT4* is specifically induced by PTI, implying distinct roles for individual LHTs in different branches of the immune response. This specificity may reflect divergent regulatory circuits controlling amino acid flux during early defense signaling versus pathogen-facilitated susceptibility. Collectively, these findings converge on a model where amino acid transporters function as central integrators of nutrition and defense. Pathogens appear to manipulate this network to create favorable conditions for infection, while plants deploy regulatory circuits, including ANAC046, to recalibrate amino acid fluxes in defense. Deciphering this regulatory network opens avenues for strategic modulation of amino acid transport and metabolism.

By employing integrative -omics approaches combined with dynamic transcriptional simulations and predictive network modeling, our study identified ANAC046 as a master regulator of amino acid metabolism during ETS **(Fig. 2** and **Fig. 4)**. This finding builds upon previous reports identifying ANAC046 as a central hub in transcriptional networks governing both developmental transitions and senescence(Oda-Yamamizo, Mitsuda et al. 2016, Huysmans, Buono et al. 2018, Mishra, Sun et al. 2018, Mahmood, Zeisler-Diehl et al. 2019). Its involvement in ETS-associated metabolic reprogramming highlights ANAC046 as a potential node of convergence where environmental stress responses and developmental pathways intersect. This multifunctionality is consistent is consistent with its broad expression profile across diverse cell types, with notably higher expression in mesophyll and vascular tissues **(Fig. 2)**, while, many amino acid-related genes, including transporters, exhibit predominant expression in mesophyll and companion cells. This cellular overlap suggests that ANAC046 may coordinate metabolic reprogramming in specific cellular contexts that are critical for both nutrient allocation and immune modulation during stress conditions. The distinct yet intersecting expression domains of ANAC046 and amino acid-related genes further support a model in which cell-type-specific transcriptional regulation fine-tunes the metabolic landscape to meet the physiological demands of both growth and defense. Previous studies have revealed a few TFs as key regulators of amino acid metabolism and transport in plants, integrating environmental signals and internal metabolic cues to fine-tune gene expression for optimal growth and stress adaptation. For example, DC3000 infection is known to activate a a basic leucine-zipper (bZIP) domain TF in Arabidopsis, which induces *UMAMIT* and *SWEET* genes that facilitate nutrient efflux from host cells—a process that can enhance pathogen proliferation(Prior, Selvanayagam et al. 2021). Specifically, bZIP11 supports pathogen replication by promoting nutrient availability; its loss impairs pathogen growth. Notably, akin to ANAC046 **(Fig. 4)**, loss of the bZIP11 leads to reduced pathogen growth(Prior, Selvanayagam et al. 2021), suggesting that bZIP11 facilitates pathogen proliferation by modulating host nutrient metabolism. Another key regulator, ANAC032, modulates primary nitrogen assimilation and amino acid catabolism by upregulating genes involved in branched-chain amino acid degradation and transport, including UMAMIT family members(Allu, Brotman et al. 2016). Our study further reveals that ANAC046 directly targets amino acid transporter genes, such as AAPs and LHTs, suggesting its broad influence over nutrient mobilization during ETS. These findings raise compelling questions about the possible coordination between bZIPs and ANAC TFs, and whether they, along with other regulators, orchestrate a broader transcriptional program governing amino acid metabolism and transport during biotic stress. Such insights will be critical for understanding how transcriptional networks integrate metabolic and immune responses, and for designing strategies to disrupt pathogen exploitation of host nutrient resources.

Our study demonstrates the power of systems biology in uncovering novel components of complex biological processes by integrating diverse “-omics” datasets. By analyzing the global network architecture derived from transcriptomic and interactomic data, we identified topologically distinct nodes—particularly hub nodes—that are significantly enriched in amino acid–related genes **(Fig. 5** and **Fig. 6)**. These central positions within the network suggest functional importance in coordinating plant metabolic and immune responses. Network centrality analyses further enabled the prioritization of seven previously uncharacterized genes as potential novel players in plant defense. This approach builds on earlier findings that topological features such as hubs, bottlenecks, and core-periphery structures can effectively predict components of the plant immune system(Mukhtar 2013, Mukhtar, McCormack et al. 2016, Mishra, Sun et al. 2017, Mishra, Sun et al. 2018, Mishra, Kumar et al. 2019, Mishra, Kumar et al. 2021). Importantly, these structural attributes also distinguish between conditionally required genes and those essential for general development or morphology(Ahmed, Howton et al. 2018). The construction of the MID*ata* represents a major advance in the application of systems biology to plant research. MID*ata* is a unified, user-friendly platform that allows researchers to explore gene function and regulatory architecture through integrative, multi-omics analysis. By incorporating network centrality measures like degree, betweenness, and closeness, it helps identify key regulatory nodes and generate biologically meaningful hypotheses. MID*ata* bridges a critical gap in plant biology by enabling systems-level insights and promoting the use of network-based approaches for functional discovery. Taken together, our findings highlight the utility of integrative, network-based approaches for functional gene discovery. By combining high-resolution omics data with mathematical modeling and computational tools, we not only identified new immune-related components but also uncovered ANAC046 as a key transcriptional regulator at the intersection of metabolic and defense pathways. This work underscores the potential of systems-level analyses to advance our understanding of plant biology and facilitate the discovery of critical regulatory nodes in complex cellular networks.

## Conclusion

In summary, comparative coexpression networks followed by topological and functional analyses identified amino acid-genes-related enrichment in the modules of the DC3000 network that exhibit increased clustering coefficients and connectivity. Dynamic transcriptional modeling and computational simulation revealed an ETS-specific TF ANAC046 as a master regulator involved in diverse biological processes including transcriptional regulation of DC3000-induced amino acid-related genes. Our plant pathology and molecular analyses illustrated that ANAC046 is preferentially involved in ETS rather than PTI. Integration of transcriptome and interactome data provided insights into the topological features of amino acid-related genes as well as their involvement in plant immunity. Finally, integrative network analyses led to the discovery of additional novel players in the plant immune system. Taken together, our integrative systems biology analyses entailing generation, integration, modeling, simulation, and inference of the multi-omics data demonstrated substantial predictive power to identify novel components of plant immunity.

## Author contributions

N.K., B.M. and A.F. conducted bioinformatics analyses, B.M performed the single cell RNA-Seq data analysis, Y.S., D.T. and T.W.D. conducted wet-lab experiments and analyzed the data, K.P.-M. and M.S.M. coordinated the research program and oversaw the experimental work, N.K., B.M. and M.S.M. wrote the manuscript. All authors discussed the results, critically reviewed the manuscript and provided feedback.

## Data and materials availability

All data needed to evaluate the conclusions in the paper are present in the paper and/or the Supplementary Materials. Additional data related to this paper may be requested from the authors. The reported mutant seeds and plasmids can be provided by MSM pending scientific review and a completed material transfer agreement. The MID*ata* database can be found at https://midata.cas.uab.edu/. The weighted *k*-shell decomposition analysis Cytoscape plugin app can be downloaded at http://apps.cytoscape.org/apps/wkshelldecomposition.

## Materials and Methods

### Plant material and growth conditions

T-DNA insertion lines were obtained from ABRC for SALK_025328C (AT2G36720), CS860541 (AT2G01210), SAIL_900_B07 (AT5G63930), SALK_081193C (AT1G72180), SALK_109214C (AT1G34420), SALK_205581C (AT5G08520), SALK_081049C (AT3G08710), and SALK_130637C (AT5G44700).

Genotyping primers were designed at SIGnAL’s T-DNA primer website. Homozygosity of all mutants was confirmed prior to experiments. Col-0 was used as wild type, and four-week- old soil-grown plants were used for all experiments. Plants were grown in environmentally controlled growth chambers at 21 °C with a 12/12 h light/dark photoperiod. To generate transgenic lines, Col-0 was transformed using the floral dip method*(Clough and Bent 1998)*.

### Reactive oxygen species (ROS) burst and pathogen growth measurement

ROS burst response was measured following a previously published protocol(Bisceglia, Gravino et al. 2015) with minor modifications. Briefly, after overnight water treatment of leaf discs, water was replaced with 100 l working solution containing 100nM flg22 (Genscript), 34 μg/ml luminol (Sigma) and 20 μg/ml horseradish peroxidase (Sigma). Then luminescence was detected immediately using a 96-well plate reader for 36 minutes with a signal integration time of 1 sec. Pathogen growth assay was performed as described before(Liu, Afrin et al. 2019). Briefly, *Pseudomonas syringae* pv. tomato DC3000 (hereafter DC3000) or hrcC*^-^* (a type III secretion system mutant of DC3000) with OD_600nm_=0.002 in 10mM MgCl_2_ was syringe-infiltrated into the leaves of four-week-old *Arabidopsis* plants. Six plants per genotype and four leaves per plant were infiltrated and collected for the following pathogen quantification.

### RNA extractions and RT-qPCR

*Arabidopsis* leaf samples were snap frozen in liquid nitrogen at 0 hour, 24 hours, 48 hours and 72 hours after the infiltration of DC3000 or hrcC^−^ at OD600nm=0.002 for RNA extraction. Total RNA was extracted using the RNeasy Plus Mini Kit (Qiagen) and treated with DNAse using the DNA- free kit (Ambion). RNA integrity was verified on an agarose gel. cDNA was synthesized using Super Script IV reverse transcriptase first-strand synthesis kit (Invitrogen), using 2 µg RNA in a 10 μL reaction. RT-qPCR was performed using 2X PowerUp SYBR green master mix (Applied Biosystems, ThermoFisher Scientific) on ABI 7500 Fast PCR System (ThermoFisher Scientific) with the following settings: 50°C for 2 min and 95°C for 10 min followed by 40 cycles of 95°C for 15 sec, 55°C for 15 sec and 72°C for 1 min.

## Bioinformatics and statistical analyses

The detailed bioinformatics and statistical analyses are mentioned in supplementary text. **Coexpression networks construction and analysis:** The weighted gene correlation network analysis (WGCNA) was conducted with the genes expressed in *Pseudomonas syringae* pv. tomato DC3000 (hereafter DC3000) and DC3000*hrpA*^−^(hereafter hrpA^−^)(Langfelder and Horvath 2008). The soft threshold 14 and 18 for DC3000 and hrpA^−^ respectively were used and networkType was set as ‘signed’ to construct the topological overlap matrix (TOM). The hierarchical clustering of the genes based on the TOM dissimilarity measure was done using the default parameter and flashClust function(Langfelder and Horvath 2008). The module identification was done using cutreeDynamic function with minClusterSize of 30. The visualization of the network so obtained was done using the open-source software platform Cytoscape (version 3.8.0)(Shannon, Markiel et al. 2003).

### Gene ontology (GO) analysis

Functional enrichment analysis of genes or proteins in coexpression networks, interactomes, and gene regulatory networks was achieved by TAIR GO, Enricher, ClueGO, Metascape, and DAVID with a significant threshold (*p*<0.05)(Bindea, Mlecnik et al. 2009, Jiao, Sherman et al. 2012, Kuleshov, Diaz et al. 2019, Zhou, Zhou et al. 2019).

### Dynamic regulatory event mining

The Scalable Models for the Analysis of Regulation from Time Series (SMARTS)(Wise and Bar-Joseph 2015) tool was used to reconstruct condition-specific dynamic regulatory networks in an unsupervised method by utilizing time-series gene expression data and static gene regulatory network to identify key regulators in the dynamic regulation pathways. The log2 fold change values of DEGs (fold change ≥ 2 and q ≤ 0.01) were calculated and plotted. The analysis included the log fold change of deferentially expressed genes identified in the DC3000 and hrpA^−^ time series data. The y-axis represents log2 normalized expression and x-axis represents the time points of expression data. The color of lines corresponds to expression-based clustered genes involved in certain biological processes. The dynamics of transcription factors (TFs)-activated pathways were generated by the TAIR GO ontology function.

### Single cell RNA-Seq Data Analysis

Two latest single cell RNA-Seq data sets (GSE226826 and GSE213625) were downloaded from NCBI GEO for the analysis(Zhu, Lolle et al. 2023, Nobori, Monell et al. 2025). ScRNA-Seq and multi-ome data analysis was performed using Cell Ranger, Seurat, and Signac workflow. First, Cell Ranger was utilized for the alignment of raw sequencing reads to the reference genome(Zheng, Terry et al. 2017). Next, Seurat and Signac workflow was be used for data preprocessing and quality control to filter out low-quality cells(Satija, Farrell et al. 2015, Stuart, Srivastava et al. 2021). Normalization was performed followed by the identification of highly variable features. The data was scaled, integrated with harmony, and dimensionality reduced by UMAP, and clustering was performed using graph-based methods(Hao, Hao et al. 2021). Differential expression analysis is conducted to identify marker genes for each cluster. The cluster annotation was performed by marker genes. The amino acid module scoring was performed by UCell module scoring methods(Andreatta and Carmona 2021).

### Network centrality analyses

NetworkX and Cytoscape network analysis tools are used to find the network topological indices for each node. We implied network topology analysis for the degree, betweenness, cluster coefficient, connectivity, and weighted *k*-shell decomposition centralities. In an undirected network, the degree of a node is the number of connections or the edges connected to it. In a complete network/graph all nodes (*v*) of a network (G) must have a minimum of 1 edge. Betweenness centrality is based on the shortest path of the network. The shortest path in a complete network (G) is the minimum distance between two given nodes (s and t). The betweenness centrality (C_B_) a node (*v*) is the count of shortest paths that pass through that node.

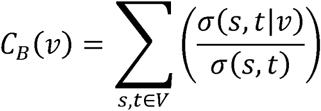

Where, ν is the set of nodes, σ(*s,t*) is the shortest (s, t) path between *s* and *t* nodes. The clustering coefficient (*C_u_*) is a measure of the degree to which nodes in a graph tend to group to form a cluster.

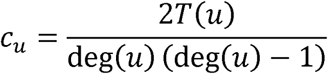

where T(u) is the number of triangles through node u and deg(u) is the degree of u.

Node connectivity (NC_(v)_) is the average connectivity of neighbors of the given vertex (v) within the network(G).

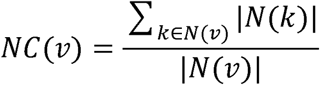

Where, N(*v*) is a set of neighbors of vertex *v*.

### Weighted *k*-shell decomposition

The Weighted k-shell decomposition is a method that ranks the most important nodes in a network and partitions them into shells based on that rank. This algorithm assigns a weight to nodes based on the degree of that node and the adjacent nodes. The Weighted k-shell-decomposition method is implemented as a Cytoscape App, *<u>wk-shell-decomposition</u>* (version 1.1.0). This app is widely used with > 2,500 downloads (April 19, 2025). The app is developed using the Apache Maven software project management and comprehension tool for Java projects. The Java SE Development Kit 11 and Maven 3 is used for the App. This app will generate a column “_wkshell” on the network’s node table that stores the rank and partitions the network into ordinal k-shells. The app then adjusts the layout of the network to packed concentric rings sorted by k-shell and applies a gradient to the color of the nodes by k-shell. The nodes in the inner shells are given red color and nodes in the outermost shell are assigned blue color. This app can be run by accessing it under the apps menu or by calling the command: “wkshell decompose”. The github repo of the app is: https://github.com/finnor/wk-shell-decomposition.

### Multi-omics Integrated Database of *Arabidopsis thaliana* (MID*ata*) database

The <u>MID*ata*</u> database is the collection of a wide variety of networks collected and curated from multiple resources and generated in-house. The MID*ata* has three main components (a) the multi-omics interaction search engine, (b) network centrality (Degree, Betweenness centrality, Closeness centrality), and (c) the raw network download link. At present only *Arabidopsis* Genome Initiative identifiers (AGI IDs) are supported as queries. Four types of -omics data are available on MID*ata*: coexpression, TF-target, Phenotype, and Protein-Protein Interaction (PPI) network (PPI-experimental and PPI-predicted). AGI IDs can be used to make a query separately from each network type and, on the results page, along with a collapsible table, there is a download option as an excel sheet.

