## Supplemental Text for "Integrative Omics and Network Biology Reveal Transcriptional Changes of Amino Acid Transport in Arabidopsis Susceptibility to *Pseudomonas syringae*"

**Title:** Integrative Omics and Network Biology Reveal Transcriptional Reprogramming Underlying Disease Susceptibility in Arabidopsis–Bacterial Interactions

**Short title:** Multi-omics in disease susceptibility

**One Sentence Summary:** Systems analyses entailing generation, integration, modeling, and simulation of diverse omics data identify novel components of plant immunity.

**Authors:**

Bharat Mishra^1§†^, Nilesh Kumar^1φ†^, Yali Sun^1‡†^, Thomas W. Detchemendy^1ι^, Doni Thingujam^1,2^, Adrian Flannery^4ϒ^, Karolina M. Pajerowska-Mukhtar^1,2^, M. Shahid Mukhtar^1, 3^*

### **Affiliations:**

^1^Department of Biology, University of Alabama at Birmingham, 1300 University Boulevard, AL 35294, USA.

^2^ Department of Biological Sciences, Clemson University, 132 Long Hall, Clemson, SC 29634, USA.

^3^ Department of Genetics & Biochemistry, Clemson University, 206 Biosystems Research Complex, 105 Collings St., Clemson, SC, 29634, USA.

^4^Division of Genomic Diagnostics and Bioinformatics, Department of Pathology, WP220 West Pavilion, 619 South 19^th^ Street, University of Alabama at Birmingham, AL 35233, USA.

^†^These authors contributed equally to this work.

^§^Present address: Department of Biological Sciences, University of Notre Dame, 3028C McCourtney Hall East, Notre Dame, IN 46556 USA

^φ^Present address: IRCP: UAB Biological Data Science Core, University of Alabama at Birmingham, AL 35233, USA.

^‡^Present address: Revvity, Inc., Waltham, MA, USA

^ι^Present address: Thermo Fisher Scientific, USA

^ϒ^Present Address: Tempus AI, Inc. Chicago, IL, USA

**List of Supplementary Materials:**

**Supplementary Text**

**Supplementary fig S1 –S11**

**Supplementary Data: Table S1 – S5**

**References (35 – 68)**

**Supplementary Text**

**Multi-omics Integrated Database of *Arabidopsis thaliana* (MID*ata*) database:** The MID*ata* database is the collection of a wide variety of networks collected and curated from multiple resources and generated in-house. The MID*ata* has three main components (a) the multi-omics interaction search engine, (b) network centrality (Degree, Betweenness centrality, Closeness centrality), and (c) the raw network download link. At present only *Arabidopsis* Genome Initiative identifiers (AGI IDs) are supported as queries. Four types of -omics data are available on MID*ata*: coexpression, TF-target, Phenotype, and Protein-Protein Interaction (PPI) network (PPI-experimental and PPI-predicted). AGI IDs can be used to make a query separately from each network type and, on the results page, along with a collapsible table, there is a download option as an excel sheet. On the search page, in addition to the network analysis page, checkboxes are provided to make a selective query possible, and if ID(s) are found in multiple resources, those sources will appear as badge-pill. The database is built over MySQL (version 5.7), Apache/2.4.25 Linux server, and the front end is built over PHP (version 7) farm work. Moreover, icons are taken from Bootstrap (version 4.0) and font Awesome is used for styling and free icons.

The multi-omics data distribution and overlap are drawn using the UpsetR R package. The four data type is taken for comparison, gene-phenotype network, PPI, coexpression (COEXP) and TF-target network, only nodes is used for intersection analysis. On the left panel, horizontal bars are used to display the total numbers of proteins or genes in each data type (Phenotype, PPI, Coexpression, and TF-target) accordingly. The vertical bars are used to represent overlaps between data types. An isolated dot (red) represents unique data points for data types respectively and connected dots (blue, turquoise, or green) represent the interaction between multiple data types. The green bar represents the largest intersecting set whereas the turquoise bar represents the intersection of all data types.

The network analysis was done using NetworkX (version 2.4) Python 3 package. The degree, betweenness centrality, and closeness centrality were calculated for all networks and stored as MySQL table. For the Network Analysis tab search method, we implemented the use of MySQL query and PHP. Upon a successful query, all sources are displayed in the right panel, while in the middle panel, results are displayed as a hierarchical tree. The tree is implemented using open-source JavaScript D3 library. The tree is itself collapsible and expandable upon click. This tree has a maximum of four levels, starting from network type, experiment type (abiotic, biotic, developmental, and etc.), and the source of data. At the very end, network centralities (degree, betweenness centrality, and closeness centrality) are displayed for each successful query. In the download section, tab-separated (edges list) file link is provided for each network. The databases have been tested on Google Chrome (Version 85.0.4183.121), Microsoft Edge (Version 85.0.564.63) and Firefox (Version 81.0).

**Data curation**

MID*ata* relational database is a collection of published open sources and in-house data. The database contains 22,769 coexpressed genes with > 8.5 million interactions, 1,020 TF regulating targeting pairs, PPI data with >21,000 nodes and >1.2 million edges, and more than 6,000 genes with phenotype key terms brought together through our rigorous curation efforts for *Arabidopsis thaliana*. Further, protein-protein interaction data is divided into two groups; protein-protein interaction experimental validated (PPIE) and protein-protein interaction obtained through text mining (PPIT).

**Coexpression interaction networks:** The coexpression network (COEXP) consists of pairs that follow similar expression patterns in different biotic and abiotic stress experiments. The abiotic component covers a massive *Arabidopsis* pan and core interactions in 134 coexpression networks (*21*). While the biotic stress- responsive coexpression networks are constructed utilizing WGCNA in Arabidopsis immune pathosystem (*18*), 16 phytohormones *Arabidopsis* immune signaling networks (*41*), and Arabidopsis developmental (leaf senescence) (*42*). All biotic coexpression interactions are in- house data; thus, listed as MukhtarLab. Data from MukhtarLab consist of more than 6 million edges whereas there are 2 million edges in pan and core data and there is only 3.2 % overlap between them.

**PPI networks:** The PPI network in MID*ata* has been divided into two separate sections, PPIT and PPIE. PPIT consists of PPI, which is either predicted or found via text mining as defined in STRING (*43*), STRING v11, and PAIR V5 (*43-46*). There are 20,329 proteins with 1,266,807 interactions. Moreover, more than 1.8 million and 306,474 proteins-interacting edges are uniquely present in STRING and PAIR, respectively. Whereas the overlap between STRING (Experimental) and PAIR edges is only about 1.2%. Similarly, the PPIE category has a collection of open-source experimentally validated protein-protein interaction datasets. We curated data from STRING (with experimental evidence), *Arabidopsis* interactome map (AI-1MAIN) (*46*), plant-pathogen immune network (PPIN-1 and 2) (*47*), cell surface interactome (CSI) (*11*), literature-curated interactions (LCI) (*48*), membrane-linked Interactome Database version 1 (MIND1) (*49*) and BioGRID (*50*).

There are 24,674, 12,027, 4,071, 1,821, 1,375, and 555 edges that are uniquely present in BioGRID, MIND1, STRING (with experimental evidence), AI-1MAIN, LCI, PPIN-1 and 2 and CSI, respectively. The highest overlap, 5,453 edges, exist between AI-1MAIN and BioGRID (orange), whereas the biggest overlapping group [BioGRID, STRING (with experimental evidence), AI- 1MAIN, LCI, PPIN-1 and PPIN-2] (deep blue) with overlapping of only 5 interactions/edges.

**TF and target networks:** We curated TF and target gene regulatory network by including vast datasets from *Arabidopsis thaliana* Regulatory Network (AtRegNet) (*51*), Plant Cistrome Database (DAP_seq) (*52*), *Arabidopsis* transcriptional regulatory map (ARTM) (*52*), Curated_1 (*53*), TF2Network (Curated_2) (*54*) and Ath (*51, 53-55*). MID*ata* includes 1,020 TFs and 33,340 target genes. It is important to note that all TFs in our databases are also the targets of other TFs; consequently, the total number of nodes in this network is exactly the same as the total number of genes, and the average incoming and outgoing connection for nodes is 209, suggesting a characteristically dense network. Among total edges (3,497,458), the highest number of shared edges is 137 between Curated_1, Curated_2, Ath, and DAP_seq (blue). 2,611,526, 462,290, 89,894, 71,800 and 12,289 are the numbers of edges uniquely present in DAP_seq, ATh, Curated_2, Curated_1, and AtRegNet respectively. The largest number of shared edges is 192,893 (~5.5%) between Ath and Dap_seq.

**Phenotypes:** The phenotype data have been curated from the study by Lloyd *et al.* (*56*) and AtPID 5.0 (*57*). Further, we added in-house manually curated phenotype data from MukhtarLab. Lloyd *et al.* enlisted 2,400 genes with a loss-of-function mutant in *Arabidopsis*. Additionally, they introduced a categorical system to describe the phenotype. Genes were categorized into prioritized groups if multiple phenotypes have been observed (essential, morphological, cellular-biochemical, and conditional). Further subcategories were also described. AtPID 5.0 contains 3,919 integrated genotype-phenotype associations. Combining all we have more than 6,000 genotype-phenotype presents in MIDa*ta*. Some phenotypes are also described as “no phenotype”, representing a negative dataset. Other than these sources phenotype information was also mined from The Arabidopsis Information Resource (TAIR) (*58*). Further, all long sentences describing phenotypes were reformatted keeping only keywords and semantics to minimize redundant information.

**Supplementary Figures**

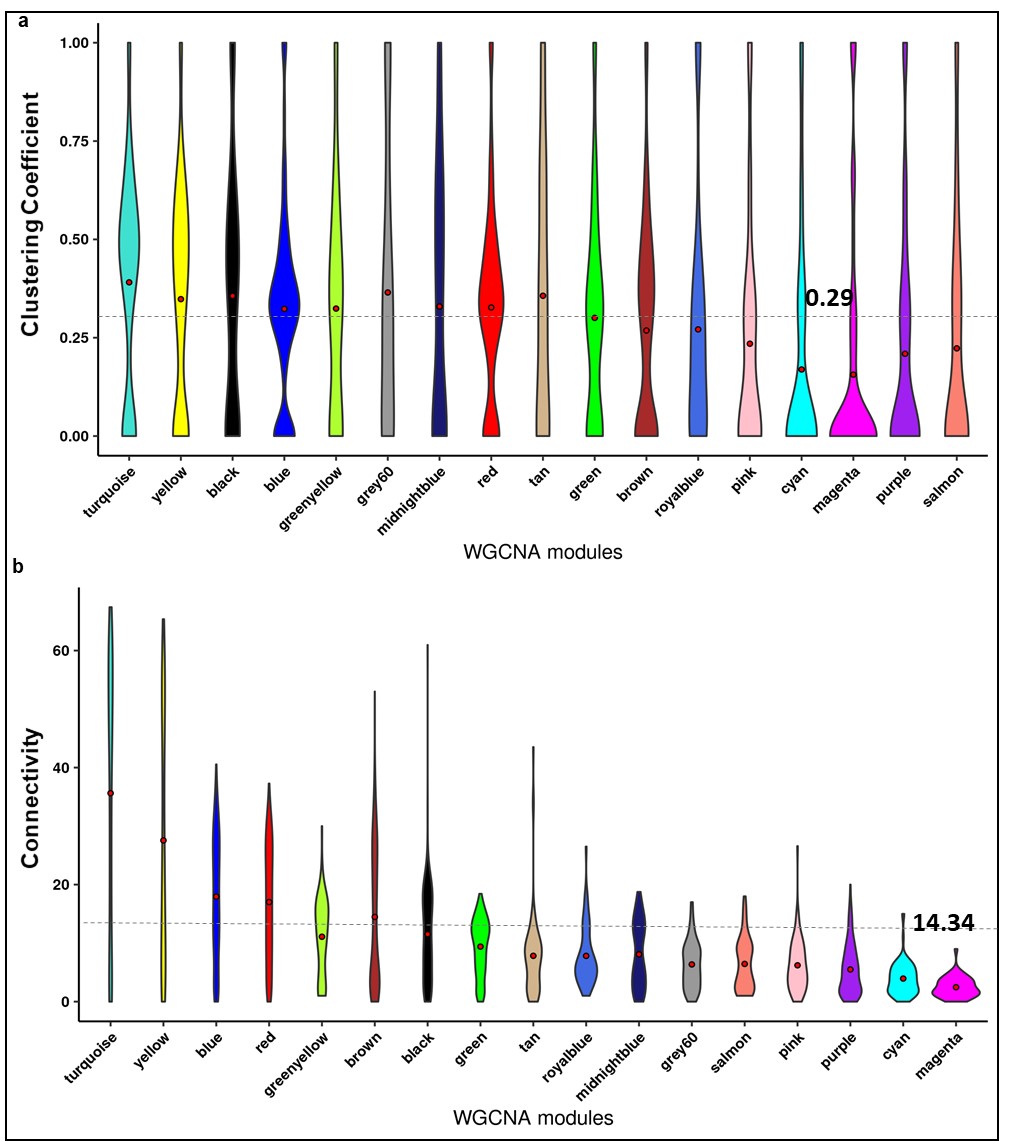

**Fig. S1:** *Pst* DC3000 coexpression network analysis. **a.** Distribution of clustering coefficient in some of the highly clustered modules in the DC3000 coexpression network. The average clustering coefficient of the entire DC3000 coexpression was considered as cutoff (0.2909475) to distinguish the highly clustered modules in the network. **b.** Distribution of connectivity in some of the highly connected modules in the DC3000 coexpression network. The average connectivity of the entire DC3000 coexpression was considered as cutoff (14.39002) to distinguish the highly clustered modules in the network.

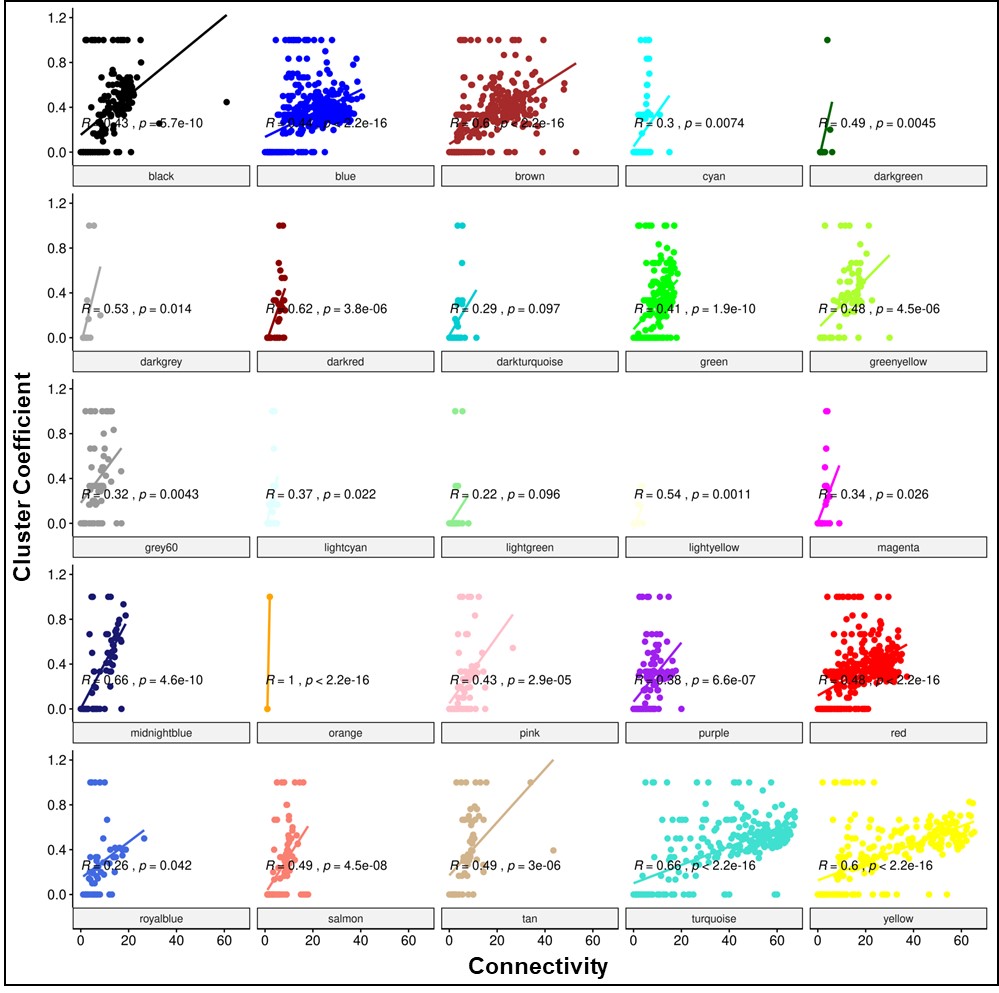

**Fig. S2:** Relationship distribution of clustering coefficient and connectivity in 25 modules in the DC3000 coexpression network. The corresponding R and *p*-value determine the most significantly clustered and connected modules in the DC3000 coexpression network.

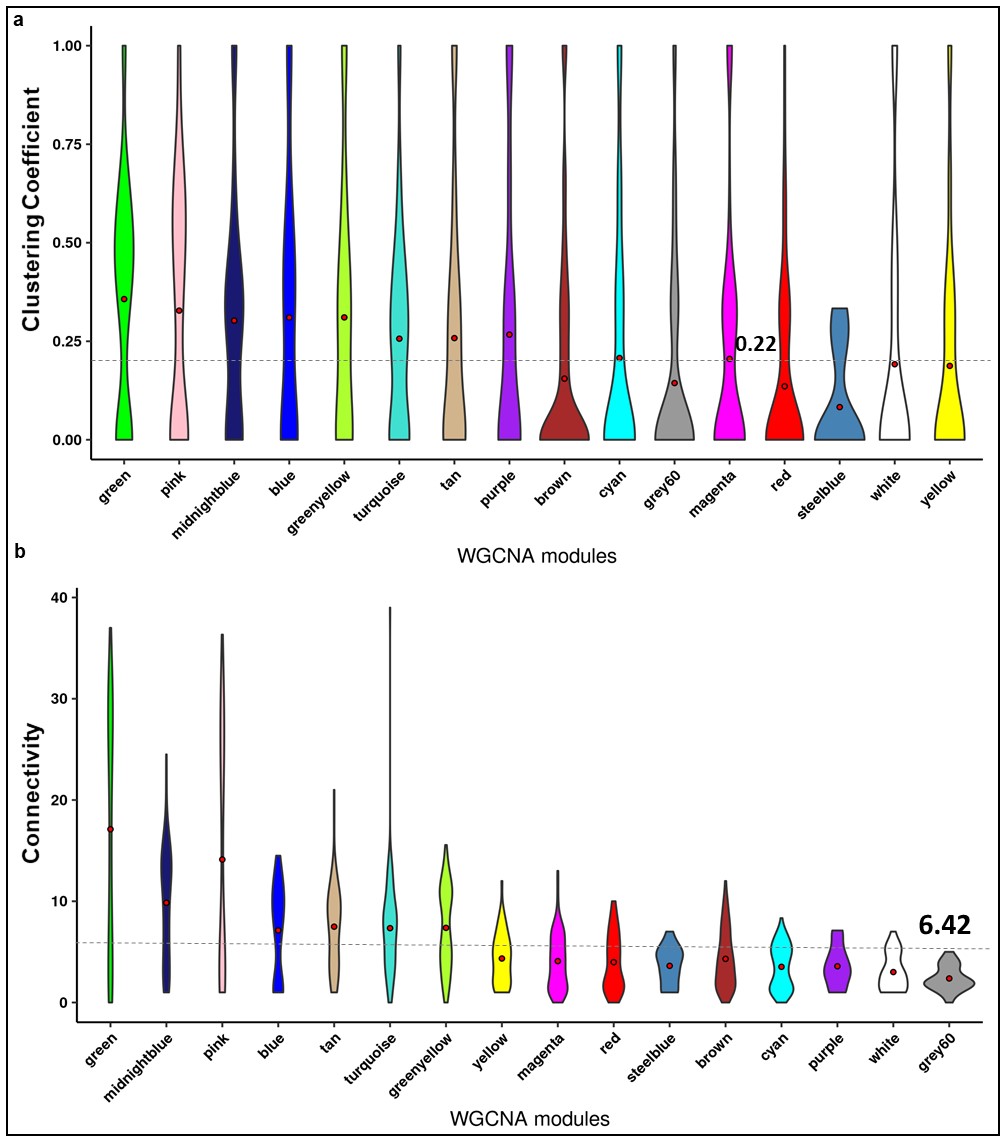

**Fig. S3:** *Pst* DC3000 hrpA^−^ coexpression network analysis. **a.** Distribution of clustering coefficient in some of the highly clustered modules in hrpA^−^ coexpression network. The average clustering coefficient of the entire hrpA^−^ coexpression was considered as cutoff (0.2177424) to distinguish the highly clustered modules in the network. **b.** Distribution of connectivity in some of the highly connected modules in hrpA^−^ coexpression network. The average connectivity of the entire hrpA^−^ coexpression was considered as cutoff (6.424122) to distinguish the highly clustered modules in the network.

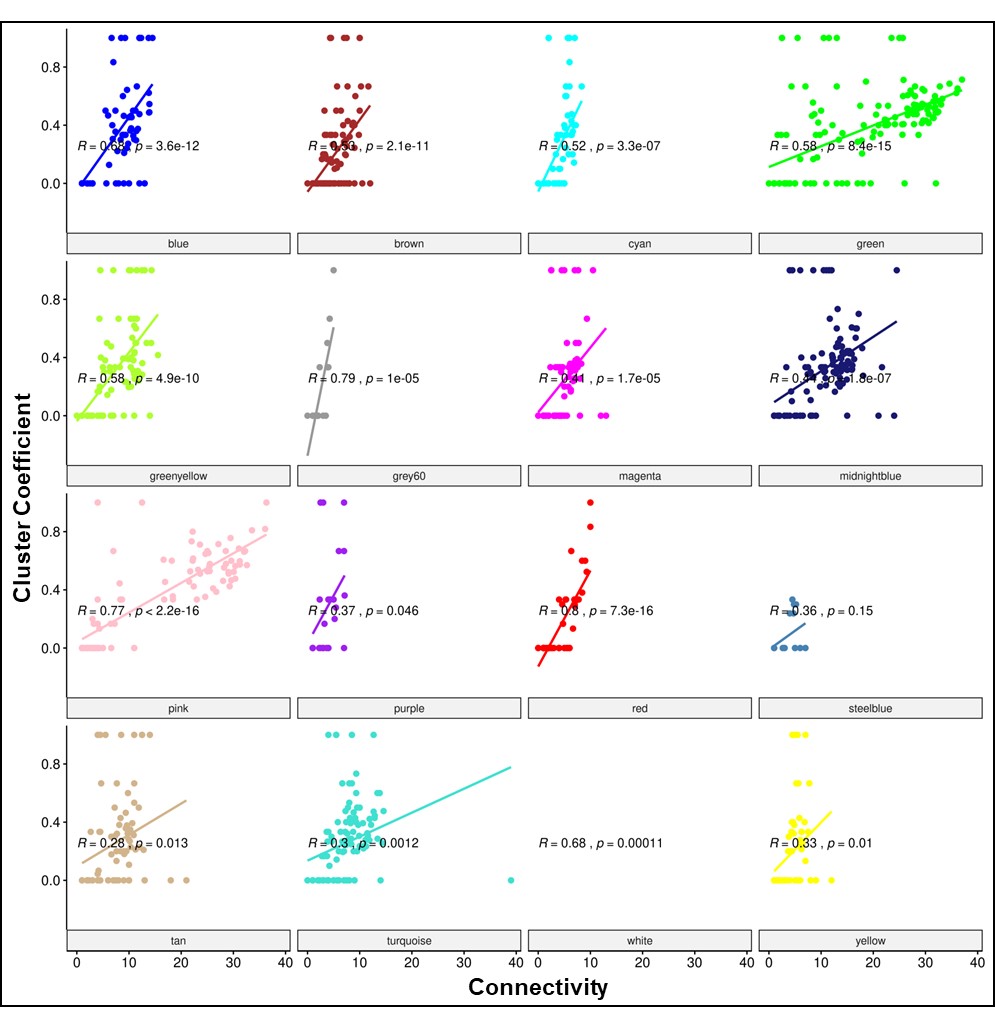

**Fig. S4:** Relationship distribution of clustering coefficient and connectivity in 16 modules in the hrpA^−^ coexpression network. The corresponding R and *p*-value determine the most significantly clustered and connected modules in hrpA^−^ coexpression network.

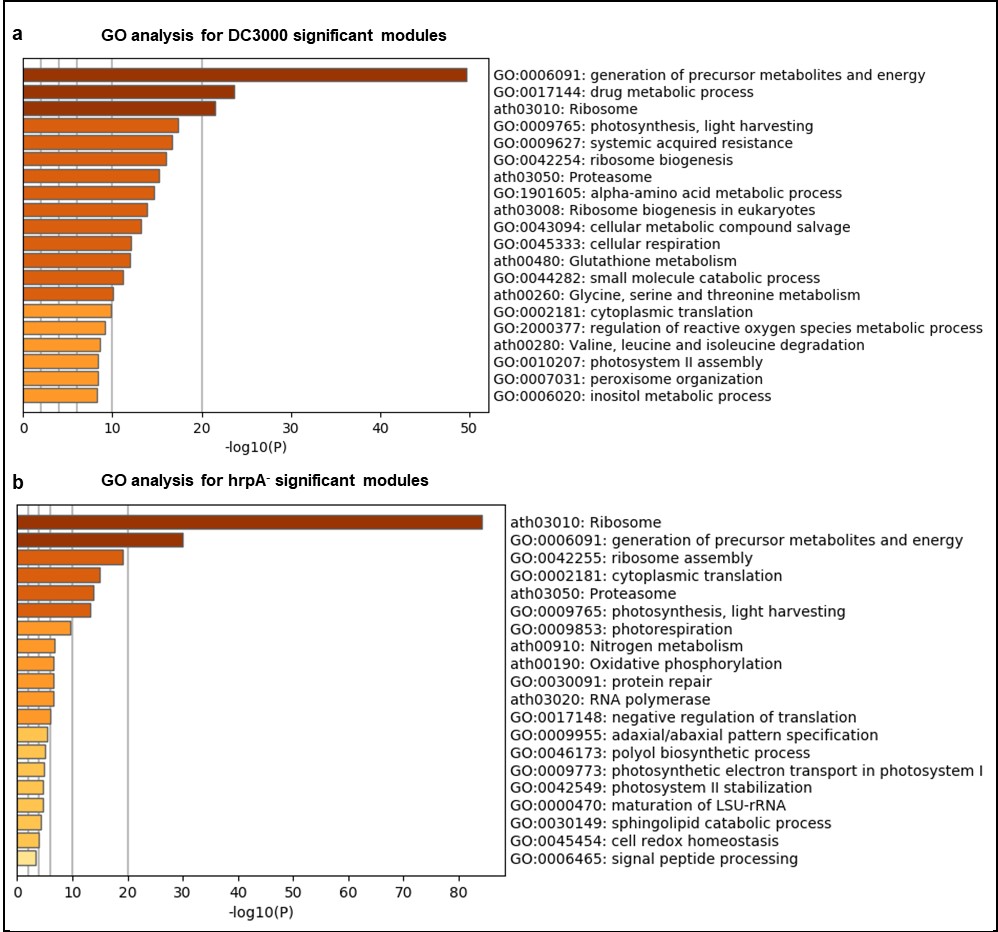

**Fig. S5:** Gene Ontology (GO) analysis of significant modules in DC3000 and hrpA^−^ coexpression networks. **a.** Some of the highly enriched pathways in DC3000 significant modules are metabolites & energy, drug metabolic process, photosynthesis, ribosome, systemic acquired resistance, proteasome, α-amino acid metabolic process, amino acid degradation process, and inositol metabolic process (*p*<0.05). **b.** Some of the highly enriched pathways in hrpA^−^ significant modules are ribosome, metabolites & energy, cytoplasmic translation, proteasome, photosynthesis, nitrogen metabolism, protein repair, sphingolipid catabolic process, and cell redox (*p*<0.05).

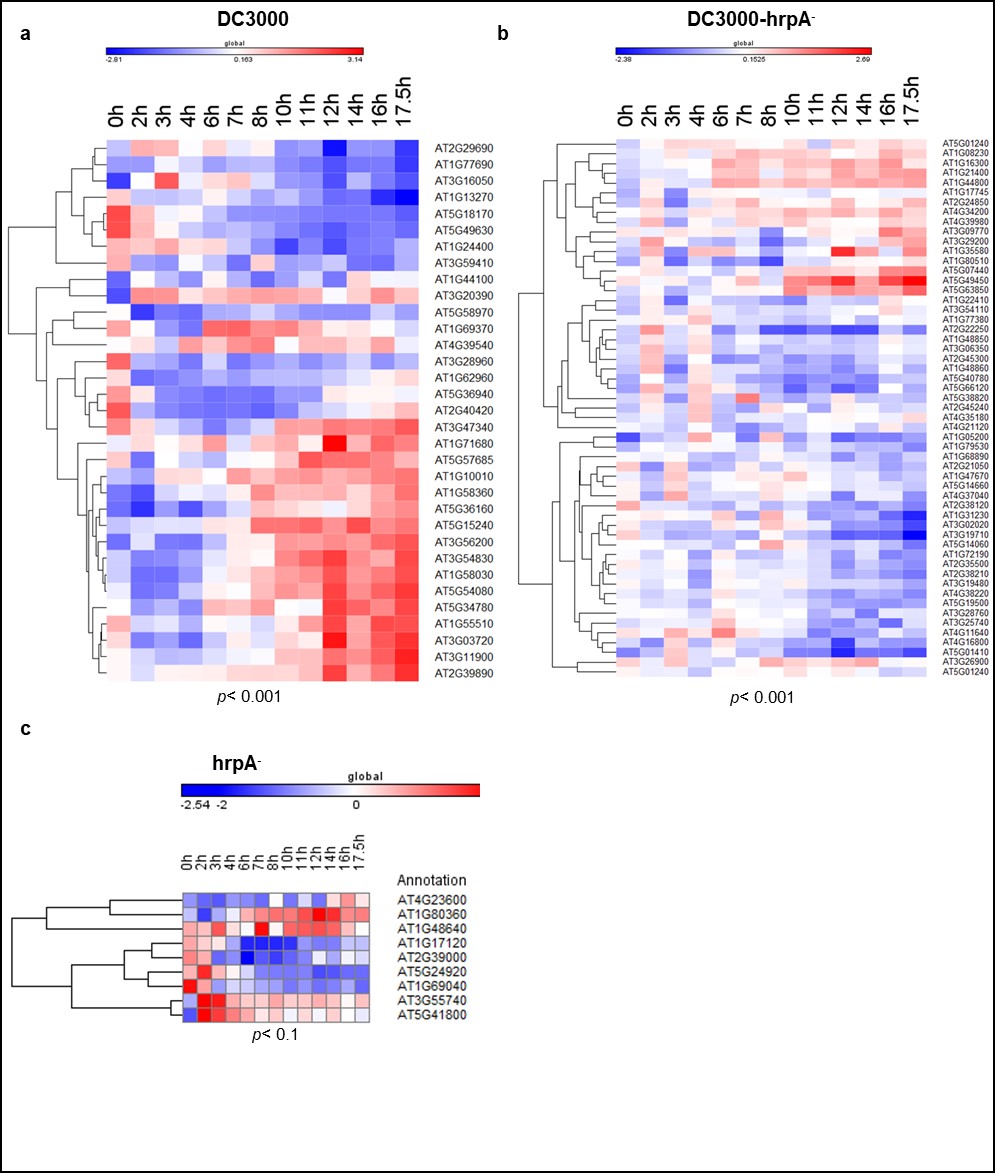

**Fig. S6:** Amino acid transporters expressed in the DC3000 and hrpA^−^ networks. **a.** Expression profiles of 33 amino acid transporters/metabolism differentially expressed genes (DEGs) in DC3000 only. **b.** Expression profiles of 55 amino acid transporters/metabolism DEGs shared between DC3000 and hrpA^−^. **c.** Expression profiles of nine amino acid transporters/metabolism DEGs in hrpA^−^ only.

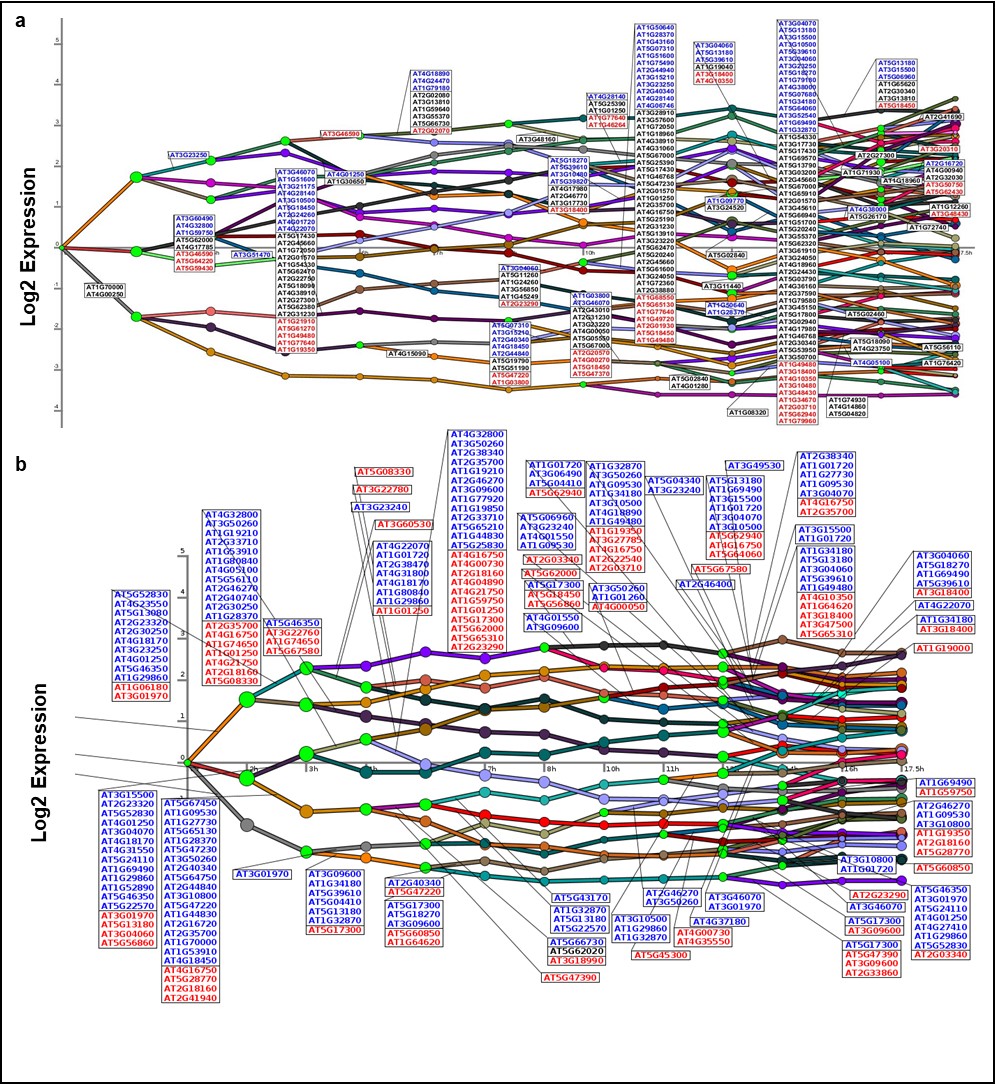

**Fig. S7:** Dynamic transcriptional regulatory event mining identifies significant transcription factors (TFs) in the DC3000 and hrpA^−^ networks. **a.** Dynamic regulatory event mining of DC3000 differentially expressed genes (DEGs) identified 198 significant TFs participate in 78 enriched paths (*p*<0.05). **b.** Dynamic regulatory event mining of hrpA^−^ DEGs identified 215 significant TFs participate in 33 enriched paths (*p*<0.05).

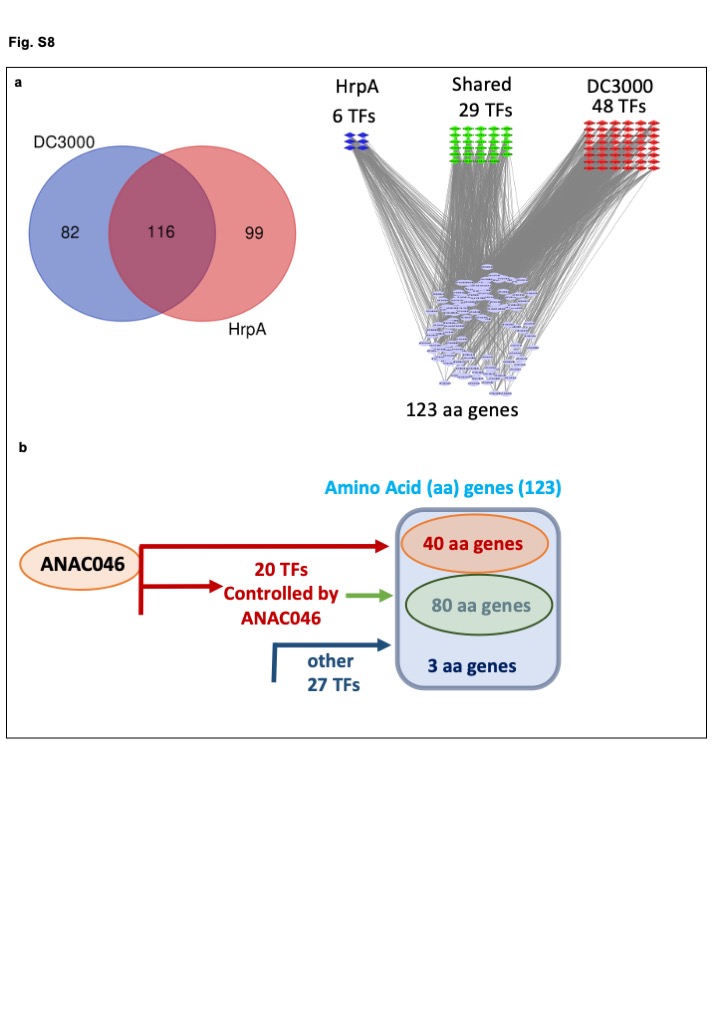

**Fig. S8: a.** Overlapping and distinct transcription factors (TFs) in the DC3000 and hrpA^−^ DEGs (left) and TFs regulating 123 amino acid genes in HrpA, DC3000-HrpA, and DC3000 (right). **b.** ANAC046 directly/indirectly regulates 123 amino acid transporter/metabolism genes and 20 out of 47 TFs, which regulate 120 amino acid transporter genes. ANAC046 direct regulatory activities are represented by light red arrows, while other TF- amino acid transporter gene interactions are represented by green and blue arrows.

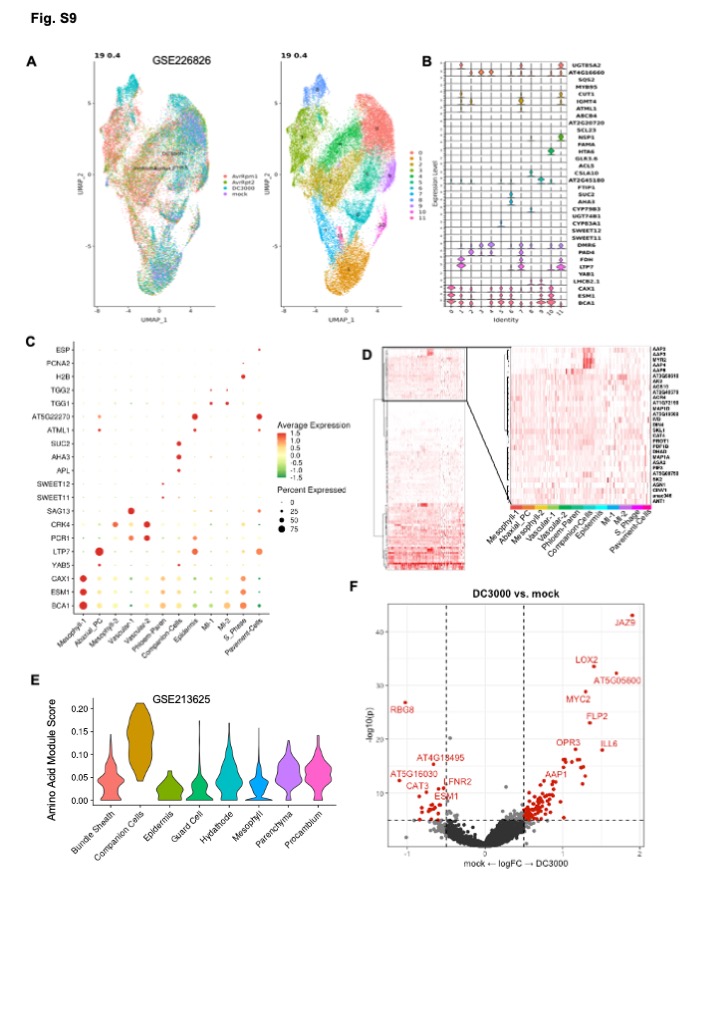

**Fig. S9:** Single cell RNA-Seq analysis of Arabidopsis leaf during pathogen infection (GSE226826). **a.** The integrated UMAP and identified cell clusters in the single cell RNA-Seq analysis at 19 dimensions and 0.4 resolutions. **b.** Violin plot of marker genes expression in individual cell clusters. **c.** Dot of marker genes expression in assigned cell types for the single cell RNA-Seq data. **d.** The Heatmap of amino acid associated genes (Fig 2A) and their interactors expression in different cell types in scRNA-Seq. **e.** The amino acid module scoring in different cell types in a different single cell dataset (GSE213625). UCell Module scoring function with genes from Figure 2A was used to perform the scoring analysis. **f.** The Volcano plot of DEGs analysis (DC3000 vs mock) in the companion cell from scRNA-Seq data.

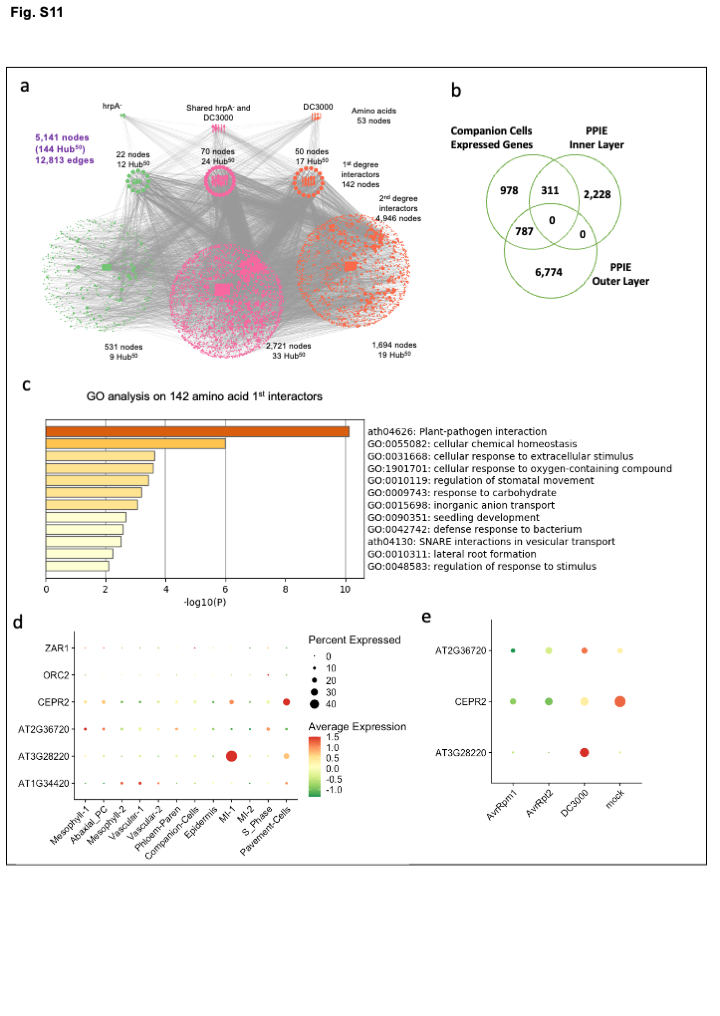

**Fig. S10:** Multi-omics data integration to identify novel players in amino acid transport. **a.** The integrated PPIE and bulk RNA-Seq differentially expressed genes in DC3000 and HrpA interactome. **b.** The overlap of DEGs (DC3000 vs mock) in Companion Cells form single cell RNA-Seq data and PPIE network. **c.** Gene ontology (GO) analysis of 142 amino acid 1^st^ degree interactors in protein-protein experimental interaction (PPIE) network identified “plant-pathogen interaction”, “stomatal movement”, “chemical homeostasis”, “response to stimulus”, “inorganic anion transport”, and “defense response to bacteria” as some of the enriched GO terms (-log10(*p*)> 2). **d-e.** Expression of selected genes in single cell RNA-seq Data in all cell types and in Companion cells.

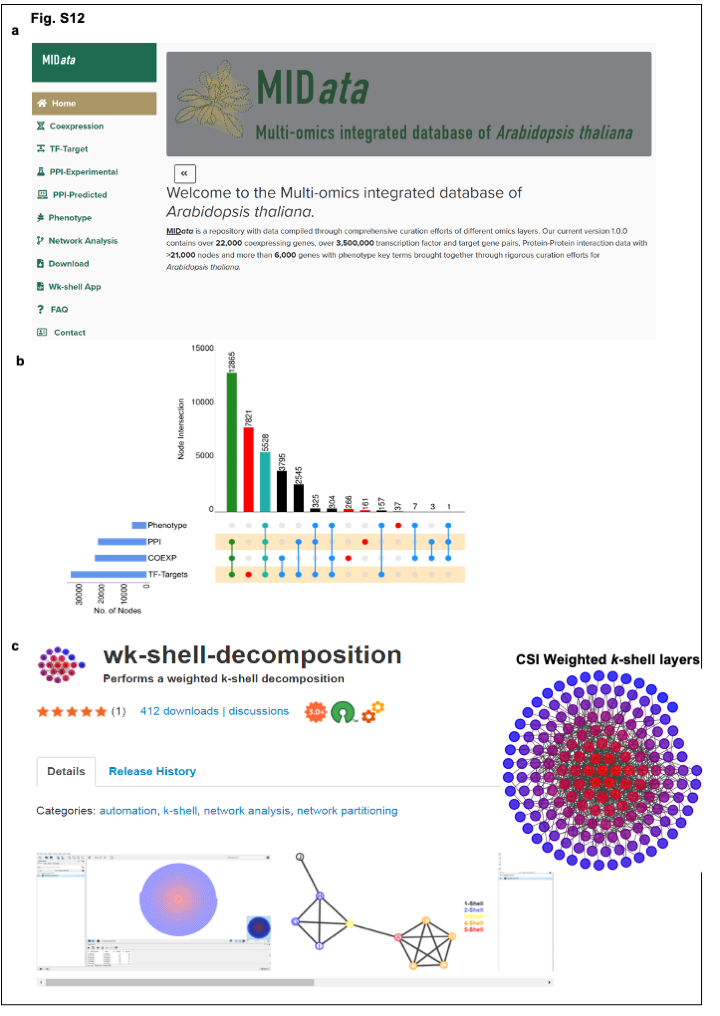

**Fig. S11:** New open-access network analyses and data acquisition tool MIData. **a.** The home page of Multi-omics Integrated Database of *Arabidopsis thaliana* (MIData). **b.** Multi-omics data acquisition and integration in MID*ata* database. (PPI: protein-protein interactions, COEXP: gene coexpression networks, transcription factor (TF)-Targets: TF gene regulatory network, and Phenotype: genetic-phenotype associations). **c.** The open-access/for public use Cytoscape plugin of weighted *k*-shell decomposition. The representative image of *Arabidopsis* cell surface interactome (CSI) weighted *k*-shell layers. Red nodes are present in the inner layer while blue and purple nodes are localized to the outer/peripheral layer of the network.

**Supplementary Data**

| **Table Number** | **Description** | **Sheet number** | **Sheet Description** |
| --- | --- | --- | --- |
| **table S1** | table S1: DC3000 and DC3000HrpA Network analysis | 1 | DC3000 Network |
|  |  | 2 | DC3000 Network Module Clusters (WGCNA) |
|  |  | 3 | DC3000 Network Centrality Analysis |
|  |  | 4 | DC3000-HrpA Network |
|  |  | 5 | DC3000-HrpA Network Module Clusters (WGCNA) |
|  |  | 6 | DC3000-HrpA Network Centrality Analysis |
|  |  | 7 | DC3000 Module Vs DC3000-HrpA Module |
|  |  | 8 | DC3000 Network Vs Random Network |
|  |  | 9 | DC3000-HrpA Network Vs Random Network |
| **table S2** | table S2. AA text mininig; DC3000 and DC3000-HrpA regulation | 1 | TAIR text mining |
|  |  | 2 | GO Biological process |
|  |  | 3 | 165 selected genes |
|  |  | 4 | 123 amino acid genes expressed in DC3000 and HrpA |
|  |  | 5 | DC3000 expression |
|  |  | 6 | DC3000-HrpA expression |
|  |  | 7 | DC3000 vs DC3000-HrpA expression |
|  |  | 8 | DC3000 Significant regulator |
|  |  | 9 | DC3000-HrpA Significant regulator |
|  |  | 10 | Significant regulator coexpressed |
|  |  | 11 | 82 regulators expressed in DC3000 |
|  |  | 12 | 48 regulators Regulating amino acid genes in DC3000 |
| **table S3** | ANAC046 interactions and master regulator role | 1 | ANAC046 Interactions from MiData database |
|  |  | 2 | DC3000 NAC046 |
|  |  | 3 | Hormone knockout NAC046 |
|  |  | 4 | Senescence (Developmental) NAC046 |
|  |  | 5 | ANAC046 regulated Amino acid GRNs |
|  |  | 6 | ANAC046 regulated Amino acid GRN node properties |
|  |  | 7 | ANAC046 Direct regulated amino acid genes and Tfs |
|  |  | 8 | Gene Description of ANAC046 targets |
| **table S4** | table S4: qPCR and PPIE significant genes based on network centrality analysis | 1 | qPCR primers used in the study |
|  |  | 2 | Mutants and their corresponding primers |
|  |  | 3 | Litrature curated phenotype diverse pathogens |
| **table S5** | table S5: Protein - Protein Interaction analysis | 1 | AI-1(main) |
|  |  | 2 | CSI |
|  |  | 3 | mind1 |
|  |  | 4 | PPIN1/2 |
|  |  | 5 | LCI |
|  |  | 6 | PPI experimental (PPIE) STRING-db |
|  |  | 7 | PPIE network analysis |
|  |  | 8 | PPIE selected genes (Centrality cutoff top 7.5 %) |
|  |  | 9 | Amino acid 1st and 2nd degree interactors in DC3000 and HrpA coexpression network |
